# A replicable and generalizable neuroimaging-based indicator of pain sensitivity across individuals

**DOI:** 10.1101/2024.06.08.597884

**Authors:** Li-Bo Zhang, Xue-Jing Lu, Hui-Juan Zhang, Zhao-Xing Wei, Ya-Zhuo Kong, Yi-Heng Tu, Gian Domenico Iannetti, Li Hu

## Abstract

Developing neural indicators of pain sensitivity is crucial for revealing the neural basis of individual differences in pain and advancing individualized pain treatment. To identify reliable neural indicators of pain sensitivity, we leveraged six large and diverse functional magnetic resonance imaging (fMRI) datasets (total N=1046). We found replicable and generalizable correlations between nociceptive-evoked fMRI responses and pain sensitivity for laser heat, contact heat, and mechanical pains. These fMRI responses correlated more strongly with pain sensitivity than with tactile, auditory, and visual sensitivity. Moreover, we developed a machine learning model that accurately predicted not only pain sensitivity but also pain reduction from different interventions in healthy individuals. Notably, these findings were influenced considerably by sample sizes, requiring >200 for univariate correlation analysis and >150 for multivariate machine learning modelling. Altogether, we demonstrate the validity of decoding pain sensitivity from fMRI responses, thus facilitating interpretations of subjective pain reports and promoting more mechanistically informed investigation of pain physiology.

## Introduction

Pain sensitivity varies widely across individuals^1^. The same nociceptive stimulus may be intolerably painful for someone, yet barely perceivable by another. What neural activity encodes such dramatic variability of pain sensitivity? This is a fundamental question in pain neuroscience. Understanding this is important to providing an objective and neurological indicator of pain sensitivity^2–4^, and supporting mechanistically informed investigations of pain by enabling detailed mapping between specific brain regions or pathways underlying pain. Furthermore, neural indicators of pain sensitivity may also aid in screening high-risk individuals for chronic pain and developing individualized pain treatment strategies^2,5,6^, since high pain sensitivity is a risk factor for chronic pain^7–9^.

Numerous previous studies have attempted to find neural indicators of pain sensitivity with non-invasive neuroimaging techniques like functional magnetic resonance imaging (fMRI)^10–14^. However, three key issues remain not well-addressed as of now. First and foremost, it is still subject to heated debate whether nociceptive-evoked brain activations can serve as neural indicators of pain sensitivity. Painful stimuli consistently activate a series of brain areas such as the primary somatosensory cortex (S1), secondary somatosensory cortex (S2), anterior cingulate cortex (ACC), insula, thalamus, and prefrontal cortex (PFC)^15,16^. An early small (N=17) but seminal study linked larger nociceptive-evoked blood oxygen level dependent (BOLD) responses in these areas with higher pain sensitivity^10^. Yet, a recent study with a large sample size (N=101) found that no single brain area significantly correlated with pain sensitivity^11^. With a slightly larger sample size (N=124), another recent study observed weak but significant correlations between nociceptive-evoked BOLD responses and pain sensitivity^17^. These discrepant findings may exemplify the replication crisis in empirical sciences^18–21^. One critical cause of non-replicability is sample size^22^. Recent studies suggest that a sample size of ∼100 in functional neuroimaging studies may be insufficient to reliably reveal the association between BOLD responses and behaviors at the inter-individual level^23,24^, especially when the true correlation is small^17,25^.

Another unresolved issue is whether nociceptive-evoked brain responses selectively track pain sensitivity rather than modality-independent stimulus factors^4^. Previous studies have shown that nociceptive-evoked neural responses can be evoked by nonpainful but equally salient tactile, auditory, and visual stimuli^26–29^, implying that these responses are not pain selective. However, recent studies have found some small brain regions^30,31^ and brain activity patterns^32,33^ that respond more to painful stimuli than to nonpainful stimuli, suggesting potentially preferential encoding of pain in the brain. Notably, these studies have mainly focused on whether nociceptive-evoked brain responses selectively track pain experience at the *within-individual* level. Rarely have previous studies quantitatively examined whether certain brain region or activity pattern correlates with individual differences in pain and other modalities alike at the *between-individual* level. In other word, it remains unclear whether there are selective neural indicators of individual differences in pain sensitivity.

The third issue concerns the generalizability of neural indicators of pain sensitivity. Pain can be induced by various types of nociceptive stimuli such as heat and mechanical pressure^34^. An ideal neural indicator of pain sensitivity should be generalizable in terms of pain types. However, few studies have employed more than one type of nociceptive stimuli, even though the neural processing of different types of nociceptive stimuli can differ^35^. It is thus of particular interest to develop generalizable neural indicators of pain sensitivity. Such neural indicators are less likely to be influenced by the specific peculiarities of individual pain types, instead capturing the common element shared across all pain types: the perceptual experience of pain.

To address these issues, we leveraged six ethnically and culturally diverse fMRI datasets (total N=1046) from three countries (China, US, and South Korea), where healthy participants received various types of painful stimuli: laser heat pain in Datasets 1, 2 & 5^14,36^, mechanical pain in Dataset 3^37^, contact heat pain in Datasets 4^17^ and 6^38^ (see Table S1 for a brief summary of each dataset). We aimed to answer four key questions (Figure 1A): (1) Do nociceptive-evoked BOLD responses reflect pain sensitivity across individuals? (2) If so, is this correlation selective to the pain modality? (3) Can we develop a generalizable machine learning (ML) model that accurately predicts pain sensitivity? (4) Which sample size is needed to reliably decode pain sensitivity from BOLD responses? First, we correlated nociceptive-evoked BOLD responses with pain sensitivity and tested the replicability and generalizability of the correlation with Datasets 1∼3 (N=794). Subsequently, we examined whether the correlation between BOLD responses and sensory sensitivity differs between pain and nonpain modalities with Datasets 1&2 (N=399), where participants also received nonpainful tactile, auditory, and visual stimuli (Figure 1B). Next, we built a multivariate ML model, named the Neuroimaging-based Indicator of Pain Sensitivity (NIPS) model, to predict pain sensitivity using nociceptive-evoked BOLD responses. We evaluated NIPS’s performance with Datasets 1&2, examined its generalizability across various pain types and diverse populations with three external healthy individual datasets (i.e., Datasets 3, 4&6, total N=611), and tested its ability to predict pain reduction from different interventions in healthy individuals (i.e., Datasets 4&6, N=487) and pain sensitivity in chronic pain patients (i.e., Dataset 5, N=36). Finally, we systematically examined the influence of sample sizes on the ability to detect the univariate correlation between BOLD responses and pain sensitivity with 80% power^39^ and achieve robust performance in multivariate predictive modeling.

**Figure 1.**
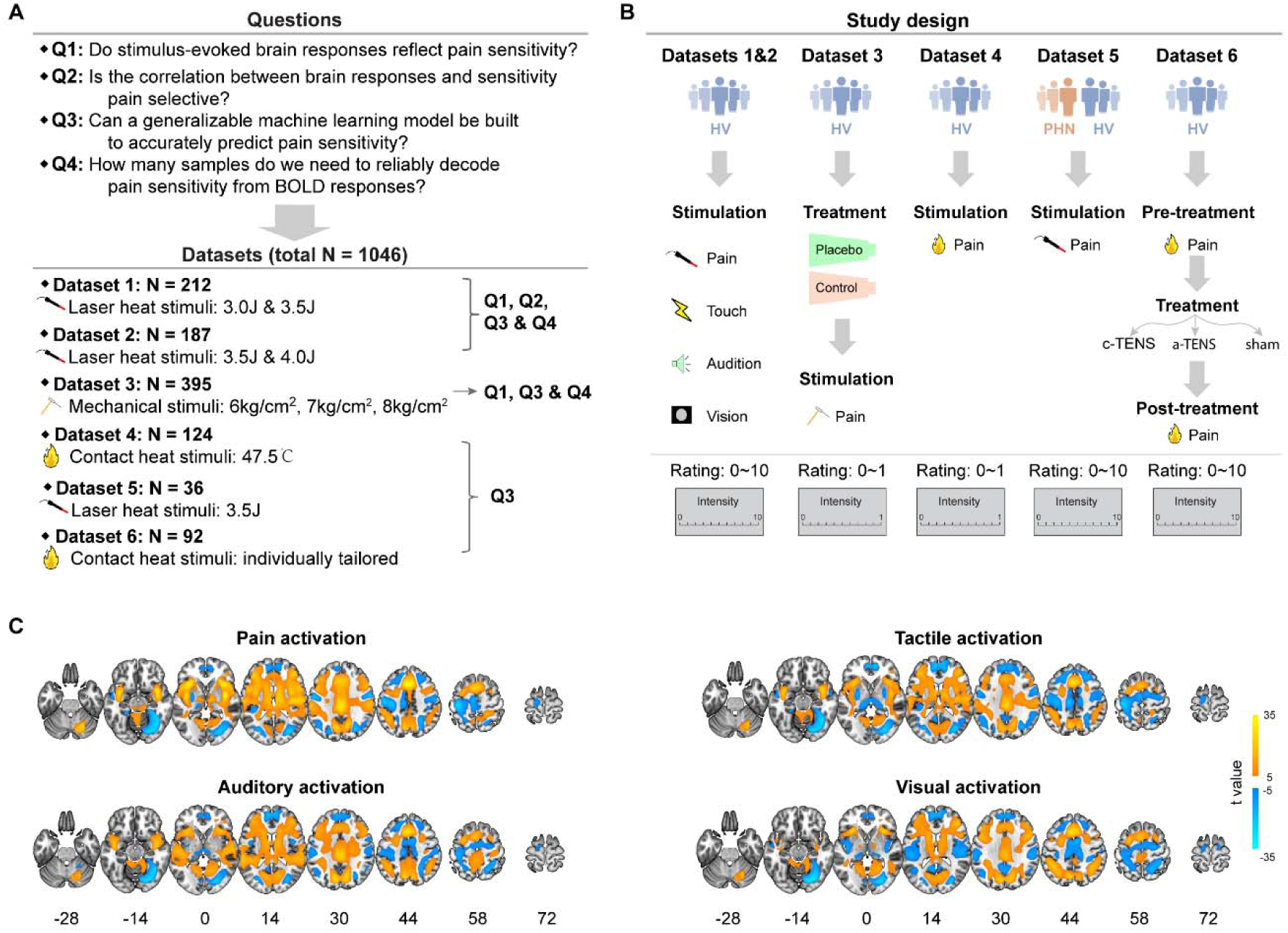
Study overview and brain activations responsive to painful and nonpainful stimuli. **(A)** Key questions and datasets used to answer these questions. Using six datasets with laser heat pain, mechanical pain, and contact heat pain stimuli, we examined the correlation between nociceptive-evoked BOLD responses and pain sensitivity, tested pain selectivity of this correlation, built generalizable machine learning models to predict pain sensitivity, and explored sample sizes needed to reliably reveal the correlation between BOLD responses and pain sensitivity. **(B)** Study design of six datasets. Datasets 1, 2, 3, 4 & 6 only recruited healthy volunteers, and Dataset 5 recruited both postherpetic neuralgia patients and healthy volunteers. In all six datasets, participants received sensory stimuli and rated the perceived intensity with a rating scale (from 0 to 10 or from 0 to 1). In Datasets 3&6, participants also received treatments (placebo in Dataset 3 and transcutaneous electrical nerve stimulation [TENS] in Dataset 6) to relieve pain. **(C)** Brain areas activated by painful and nonpainful stimuli in Datasets 1&2. Painful, tactile, auditory, and visual stimuli activated largely overlapped brain areas. HV: healthy volunteers; PHN: postherpetic neuralgia patients; c-TENS: conventional TENS; a-TENS: acupuncture-like TENS.

## Results

### BOLD responses reliably reflect pain sensitivity

We utilized Datasets 1∼3 to examine whether nociceptive-evoked BOLD responses reliably reflect inter-individual pain sensitivity. In Datasets 1&2, participants received and rated sensory stimuli of four modalities (painful, tactile, auditory, and visual) with two levels of physical intensity (low and high)^14^; in Dataset 3, participants received mechanical pain stimuli with three levels of physical intensity (6kg/cm^2^, 7kg/cm^2^, and 8kg/cm^2^) ^37^. Dataset 1 delivered laser heat pain stimuli of 3.0J and 3.5J, and Dataset 2 delivered stimuli of 3.5J and 4.0J. To make full use of the large sample size, we pooled data from the 3.5J condition (N=399) and tested the correlation between BOLD responses and pain sensitivity. We then assessed the replicability of the finding from this pooled dataset with data from the 3.0J condition in Dataset 1, the 4.0J condition in Dataset 2, and the mechanical pain data from all conditions (6kg/cm^2^, 7kg/cm^2^, and 8kg/cm^2^) in Dataset 3. Finally, we explored the effect of sample sizes on our findings by generating bootstrapped samples from Datasets 1∼2^40^. Note that Dataset 3 was originally collected to investigate placebo analgesia^37^. Here we utilized only the fMRI data in the control condition where no placebo cream was applied.

In Datasets 1&2, participants received 10 laser heat pain stimuli per condition and reported their perceptual intensity ratings for each stimulus. We quantified pain sensitivity as the mean of the 10 ratings in a condition. As in previous studies^1,11,17^, we observed considerable inter-individual variability of pain sensitivity even though the painful stimuli had identical physical intensity. Pain sensitivity ranged from 0.8 to 9.9 (mean±standard deviation [SD]: 4.54±1.72; Figure 2A and Table S4), covering almost the entire range of the 0 (‘‘no sensation’’) ∼ 10 (‘‘the strongest sensation imaginable”) rating scale. Painful stimuli also activated a wide range of typically pain-related brain areas, including the S1, S2, ACC, insula, and thalamus (Figure 1C).

**Figure 2.**
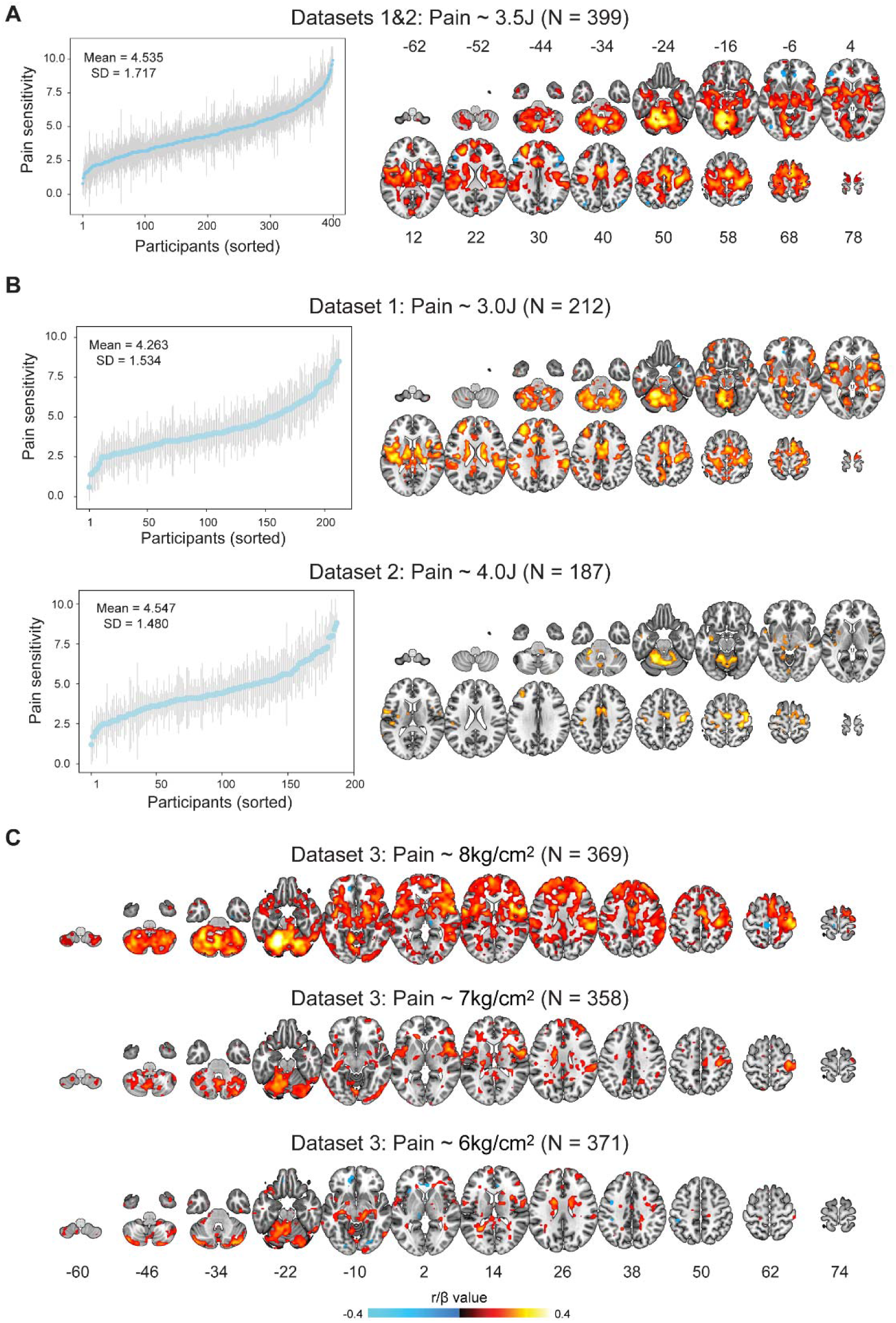
Correlations between nociceptive-evoked BOLD responses and pain sensitivity. **(A)** BOLD-pain sensitivity correlations in the 3.5J condition in Datasets 1&2. Pain sensitivity varied considerably across individuals. This interindividual variability correlated with nociceptive-evoked BOLD responses in a wide range of areas. **(B)** Replication of the correlation in other conditions in Datasets 1&2. BOLD responses correlated with pain sensitivity in the 3.0J and 4.0J conditions. **(C)** Replication of the correlation in Dataset 3. BOLD responses were associated with mechanical pain sensitivity in all conditions, demonstrating the replicability and generalizability of the correlation. Error bars in panels A&B stand for standard deviation (SD) of intensity ratings for each participant. Color bars indicate Pearson’s r values in panels A&B, and standardized coefficients (*β*) for the fixed effect of BOLD responses in mixed effects models that considered observation dependence within families in panel C.

Importantly, we found significant Pearson’s correlations between nociceptive-evoked BOLD responses and pain sensitivity across individuals (Figure 2A; p(FDR)<0.05, the same hereafter). Positive correlations were found in a wide range of areas, including but not limited to the S1, S2, ACC, insula, thalamus, precentral gyrus, supplementary motor area, precuneus, cerebellum. Negative correlations with pain sensitivity were only observed in a few areas including part of the middle frontal gyrus, inferior frontal gyrus, inferior parietal gyrus, middle temporal gyrus, angular gyrus, and inferior occipital gyrus. The absolute magnitude of negative correlations was generally smaller than those of positive correlations. To test the robustness of this finding, we also conducted Spearman’s rank correlation analysis. A similar significant correlation pattern was found (see Figure S1A for Spearman correlation map). In the analyses above, we accounted for head motion with the classical six motion parameters. To further control the potential influence of head motion, we regressed out from the BOLD signals the six motion parameters, their first derivatives, and squares of the motion parameters and first derivatives^41^. Still, the significant correlation pattern resembled that of our original finding (Figure S1B), suggesting a negligible influence of head motion on our finding.

In the foregoing analyses, we pooled the 3.5J condition in Datasets 1&2 since these datasets were assumed to only differ in pain sensitivity of participants and laser stimulation parameters based on the data collection process. Indeed, participants in Dataset 1 reported greater pain ratings to the 3.5J stimulus than those in Dataset 2 (t(397)=10.637, p<0.0001, Cohen’s d=1.067), but the sex ratio (χ^2^(1)=3.434, p=0.064) and age (t 395)=1.048, p=0.295, Cohen’s d=0.105) did not differ in these two datasets. However, there could be some unknown systematic differences between the two datasets. We therefore conducted partial correlation analysis between BOLD responses and pain sensitivity while controlled for dataset, sex, and age (see **Methods** for details). Removing the influence of these variables had some effect on the magnitude of the correlation between BOLD responses and pain sensitivity, but the general pattern of the partial correlation results remained highly similar to the original correlation results (see Figure S2A), demonstrating that dataset differences between Datasets 1 and 2 had no substantial effect on our findings.

We then examined whether our finding could be replicated with the same dataset and generalized to other datasets. In both the 3.0J condition in Dataset 1 and the 4.0J condition in Dataset 2, significant correlations were present in a series of brain areas (Figure 2B). However, fewer areas showed significant correlations in these two conditions, which could be associated with their sample sizes being only half of the pooled 3.5J condition (∼200 vs. ∼400), thereby reducing the statistical power to detect true effects. Note also that fewer areas survived correction in the 4.0J condition than in the 3.0J condition. The pain ratings were slightly but not significantly higher in the 4.0J condition than in the 3.0J condition (t(397) = 1.872, p = 0.062, Cohen’s d = 0.188), and the range of ratings was also not significantly different (4.0J condition range: 1.200 to 8.800, 3.0J condition range: 0.600 to 8.500, bootstrapping p = 0.628). These findings suggest that low correlations in the 4.0J condition were unlikely to be caused by rating differences in these two conditions, but may be related to the fact that Dataset 2 included participants with lower pain sensitivity than Dataset 1.

In Dataset 3, we also found significant associations between nociceptive-evoked BOLD responses and pain sensitivity in all three physical intensity conditions (Figure 2C). Notably, the largest effect size was consistently around 0.4 in Datasets 1∼3. Fewer areas showed significant correlations as the physical intensity decreased. One plausible explanation could be that painful stimuli of low intensity, such as 6kg/cm^2^, might not reliably induce pain sensation, potentially resulting in a poor signal-to-noise ratio of the BOLD responses. Indeed, in the 6kg/cm^2^ condition, 101 participants (27.2%) had a mean pain rating <0.014, which was defined as “barely detectable pain”. Similar to the 3.5J condition in Datasets 1&2, controlling for the influence of sex and age had no substantial effect on the results in Dataset 3 (Figure S3). Notably, in agreement with laser heat pain in Datasets 1&2 and mechanical pain in Dataset 3, a previous study analyzing Dataset 4, where contact heat pain stimuli were delivered, also observed significant correlations between BOLD responses and pain sensitivity^17^. Consistent findings in these datasets demonstrate not only the replicability of the correlation between BOLD responses and pain sensitivity, but also the generalizability of the correlation across different types of painful stimuli in ethnically and culturally diverse populations. To directly test whether classical pain-related areas encode pain sensitivity, we extracted mean signals from the S1, S2, ACC, insula, and thalamus in Datasets 1∼3 using anatomically defined masks. Significant correlations were observed in most of these regions of interest (ROI) except when the physical intensity was low (i.e., 6kg/cm^2^) in Dataset 3 (Table S5&S6).

Given the inconsistent findings in previous studies with smaller sample sizes^10,11,17^, we systematically investigated the potential influence of sample size on the BOLD responses-pain sensitivity correlation in the 3.5J condition from Datasets 1&2. Briefly, we sampled with replacement 100∼400 participants (in steps of 10) from Datasets 1&2, repeated the process 100 times, and estimated the sample size needed to detect a significant correlation in every voxel with a probability ≥0.8^39^ (see **Methods** for details). The sample size needed for different voxels varied greatly across the brain (Figure 3A). For voxels that showed significant correlations, the median sample size needed was 280, with an SD=78 (Figure 3A). To specifically focus on the pain-related ROIs, we also estimated the probability of obtaining significant correlations for the S1, S2, ACC, insula, and thalamus (Figure 3B). With α=0.01, the S2 required the smallest sample size (∼150) to reach significance with a probability of 0.8; with α=0.001, the S2 required a sample size of ∼220. These findings demonstrate that the sample size had an enormous influence on the ability to detect significant correlations between BOLD responses and pain sensitivity, and more than 200 participants might be needed to reliably detect their correlations.

**Figure 3.**
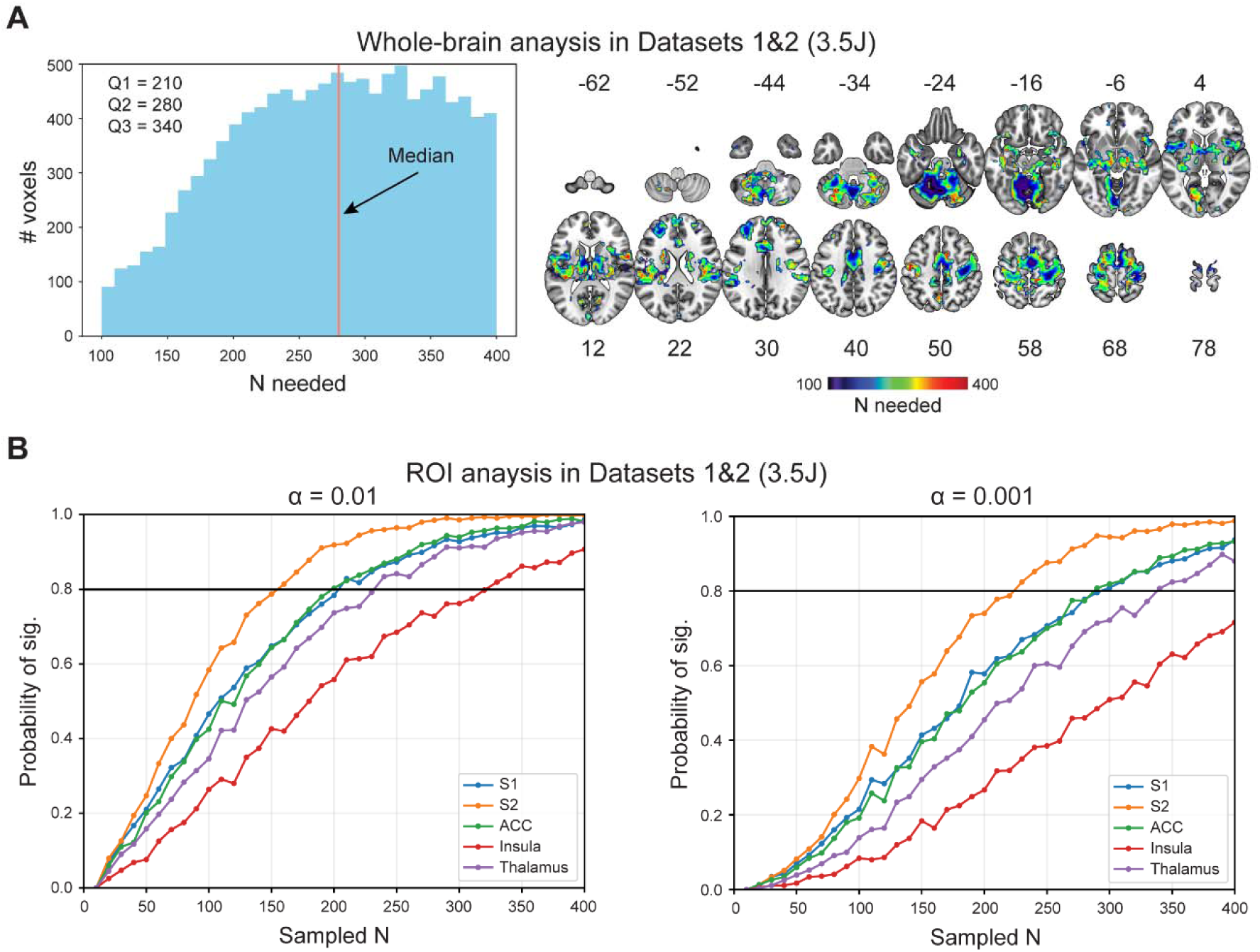
Influence of sample sizes on the detectability of the BOLD responses-pain sensitivity correlation. **(A)** Sample sizes needed to detect correlation with a probability ≥ 0.8 in whole brain analysis in Datasets 1&2. Different sample sizes were needed to reliably detect the correlations in different voxels, and a sample size of 280 was needed to detect half of the correlations. **(B)** Sample sizes needed to detect correlation in ROI analysis in Datasets 1&2. In ROI analysis, large sample sizes were still needed to reliably detect significant correlations. Q1: first quartile; Q2: second quartile; Q3: third quartile.

### BOLD responses differentially reflect pain sensitivity and nonpain sensitivity

Having shown that BOLD responses reliably reflect pain sensitivity, we then examined in the full Datasets 1&2 whether the correlation is selective to the pain modality by correlating tactile, auditory, and visual stimuli-evoked BOLD responses with the corresponding sensory sensitivity. We also compared the correlation coefficients between modalities^42^. Moreover, to rule out the potential confounding of sensitivity differences between modalities, we selected participants who had no significant sensitivity differences between modalities (see **Methods** for details) and correlated BOLD signals with sensory sensitivity in these participants with matched pain and nonpain sensitivity.

In agreement with previous findings^27^, brain regions activated by tactile, auditory, and visual stimuli were highly similar to those activated by painful stimuli (Figure 1C). However, Pearson’s correlational results exhibited distinct patterns across modalities. BOLD responses evoked by high intensity tactile stimuli correlated significantly with tactile sensitivity in several brain areas (Figure S4A), while significant correlations only occurred in some small and scattered brain areas for high intensity auditory and visual stimuli (Figure S4B&C). These results remained largely the same if Spearman’s rho was computed rather than Pearson’s r (Figure S5), or the effect of dataset, sex, and age were controlled for (Figure S2B∼D). In contrast to the high intensity condition, only a few brain areas showed significant positive correlations in the low intensity conditions for all three nonpain modalities (Figure S6). As the stimulus intensity seemed to exert substantial influences on the results, we focused on the high intensity conditions hereafter. To avoid the imager’s fallacy^43^, we then directly compared correlation coefficients between pain and nonpain modalities. For the pain vs. touch comparison, significant differences in correlation coefficients were observed in some small areas, notably in the S1, precentral gyrus, and supplementary motor area (Figure S4D). For pain vs. audition and pain vs. vision comparisons, significant differences were found in much more areas, covering typical pain-related areas, such as the S1, S2, ACC, insula, and thalamus (Figure S4D).

However, the foregoing findings might be confounded by sensitivity differences between modalities. Indeed, there were significant differences in mean ratings (i.e., sensitivity) among the four modalities (F(3,1194)=117.295, p<0.001, partial η^2^=0.228; Figure S7, Table S7). To equalize sensitivity between modalities, we adopted a within-individual sensitivity matching approach, selecting participants whose 95% confidence interval of the mean difference between pain and nonpain sensitivity contained 0^32^. With this approach, we selected 192 participants for the pain-touch matching, 171 participants for the pain-audition matching, and 147 participants for the pain-vision matching. In the matched samples for pain-touch and pain-audition, we found that nociceptive-evoked BOLD signals correlated with pain sensitivity in numerous areas (Figure 4A&B). In the matched samples for pain-vision, we observed significant correlations with pain sensitivity in several small areas like the thalamus and supplementary motor area (Figure 4C). By contrast, in all three matched samples, BOLD responses in few, if any, areas showed significant correlations with tactile, auditory, and visual sensitivity (Figure 4A∼C). Directly comparing correlation coefficients between modalities in these matched samples showed a diverse array of regions had greater correlations with pain sensitivity than with nonpain sensitivity (Figure 4D). Altogether, these findings demonstrate that there are quantitative differences in the correlation between BOLD responses and pain vs. nonpain sensitivity.

**Figure 4.**
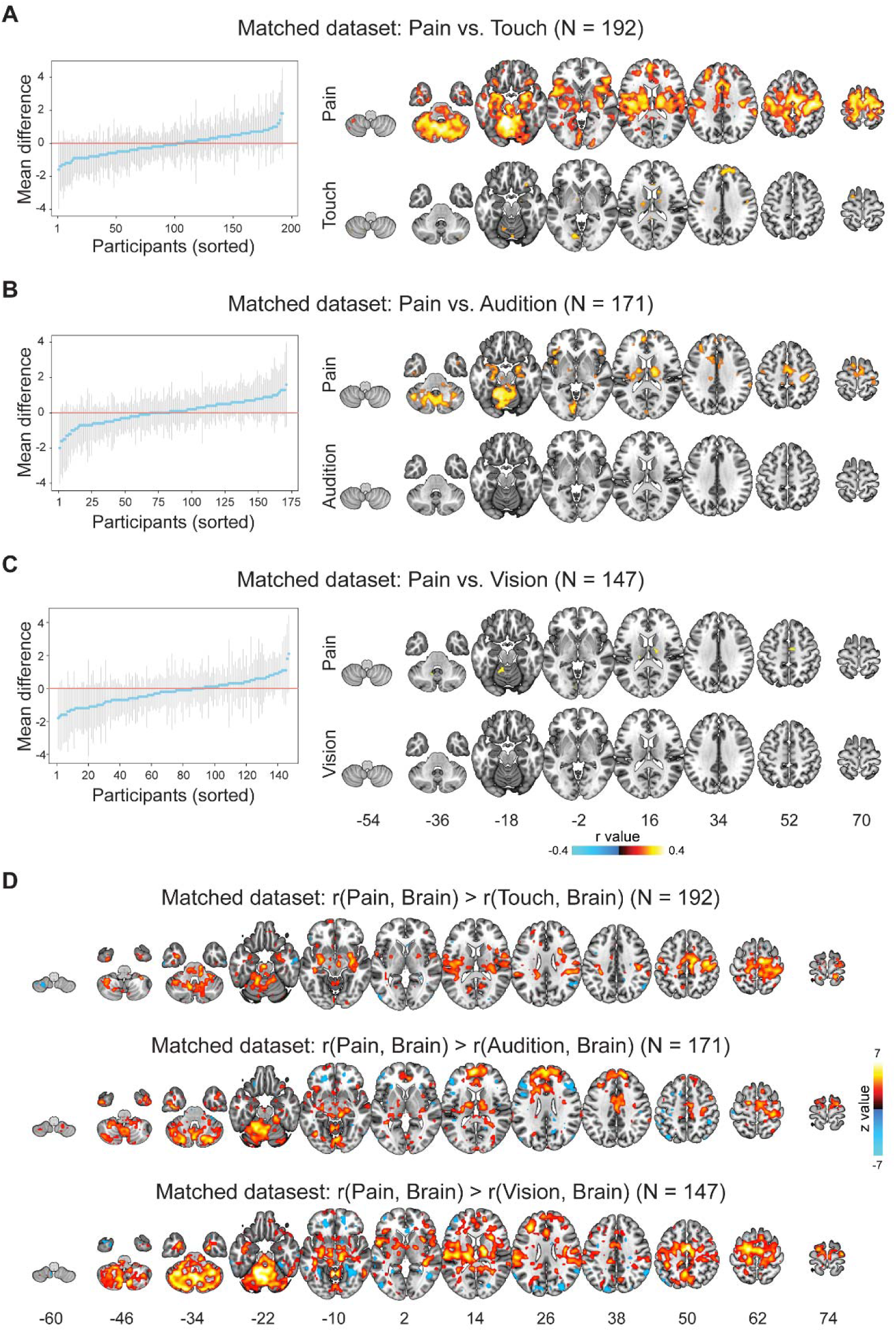
Correlation between BOLD responses and sensory sensitivity after sensitivity matching. **(A)∼(C)** BOLD responses-sensory sensitivity correlation for pain and nonpain after sensitivity matching. In sensitivity matching, we selected participants whose mean pain and tactile (A), auditory (B), or visual (C) ratings did not differ significantly. After the pain-touch and pain-audition matching, nociceptive-evoked BOLD responses still correlated with pain sensitivity. After pain-vision matching, only a few areas showed a correlation with pain sensitivity. In contrast, far fewer areas correlated with tactile/auditory/visual sensitivity in all matchings. **(D)** Comparisons of BOLD-sensitivity correlation between pain and nonpain modalities after sensitivity matching. Direct comparisons of correlation coefficients showed larger correlations for pain than for touch, audition, and vision. Error bars in (A)∼(C) stand for 95% confidence intervals for the rating differences for each participant.

### NIPS accurately predicts sensitivity to various pains in diverse populations

We next developed a neuroimaging-based ML model (i.e., NIPS) to predict pain sensitivity across individuals. We first split randomly the pooled 3.5J data in Datasets 1&2 into a Discovery Set (N=199) and a Holdout Test Set (N=200). NIPS was developed with data in the Discovery Set and tested in the Holdout Test Set. To ensure NIPS’s generalizability, we assessed its predictive power in external Dataset 3 (N=395) and Dataset 4 (N=124) with healthy participants, and Dataset 5 (N=36) with postherpetic neuralgia patients. As in the univariate analyses, we also examined the effect of sample sizes on NIPS’s performance.

To build the model, we used the least absolute shrinkage and selection operator-based principal component regression (LASSO-PCR; Figure 5A). In the Discovery Set, NIPS accurately predicted pain sensitivity with its 5-fold cross-validated performance as follows: r=0.531, p<0.0001, R^2^=0.242 (Figure 5B). NIPS also performed well in the Holdout Test Set: r=0.496, p<0.0001, R^2^=0.237 (Figure 5B). NIPS thus could explain >20% of the variance of pain sensitivity in both the Discovery Set and Holdout Test Set. Even though NIPS was built based on the data from the 3.5J condition in the Discovery Set, it was also able to predict pain sensitivity in the 3.0J condition (N=101; r=0.404, p<0.0001, R^2^=0.146) and the 4.0J condition (N=99; r=0.489, p<0.0001, R^2^=0.229) in the Holdout Test Set. Altogether, these findings demonstrate that NIPS can accurately predict pain sensitivity, explaining more than 20% of the variance of pain sensitivity across individuals.

**Figure 5.**
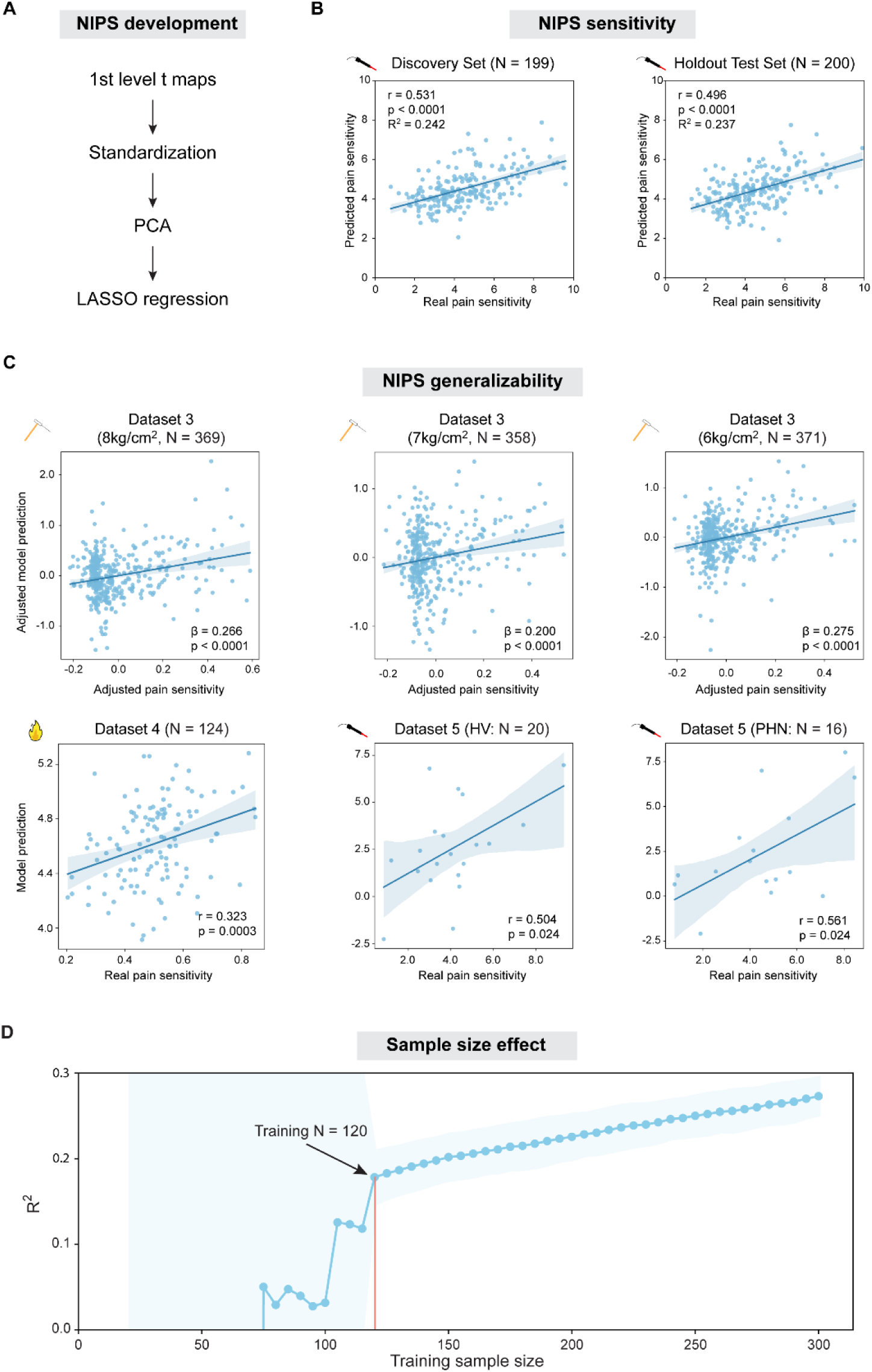
Development of NIPS and its performance. **(A)** Development of NIPS. We used first level t maps for nociceptive stimuli as features and built our model (NIPS) with LASSO-PCR. **(B)** Performance of NIPS in the Discovery Set and Holdout Test Set. In both datasets, NIPS showed good performance, explaining >20% of the variance of pain sensitivity. **(C)** Generalizability of NIPS. Although developed with laser heat pain data, NIPS significantly predicted mechanical and contact heat pain sensitivity in external datasets. Furthermore, NIPS also predicted pain sensitivity in postherpetic neuralgia patients in Dataset 5. In Dataset 3 results, “adjusted” means that random effects of family were adjusted. β is the standardized coefficient in mixed effects models. Note that real and predicted values in other datasets were not adjusted since dependent observations were not present in these datasets. **(D)** Effect of sample size on NIPS performance. NIPS’s performance varied dramatically with small training sample sizes, but stabilized and continuously improved as training sample sizes exceeded 120. Shaded regions in (B) & (C) stand for 95% confidence intervals, and shaded regions in (D) represent standard deviation (SD) of R^2^. HV: healthy volunteers; PHN: postherpetic neuralgia patients.

NIPS was developed using BOLD responses evoked by laser heat pain. To examine its generalizability, we then used it to predict mechanical pain sensitivity in Dataset 3 and contact heat pain sensitivity in Dataset 4. In Dataset 3, NIPS significantly predicted mechanical pain sensitivity in all three conditions: (1) the 8kg/cm^2^ condition: β=0.266, p<0.0001; (2) the 7kg/cm^2^ condition: β=0.200, p<0.0001; (3) the 6kg/cm^2^ condition: β=0.275, p<0.0001 (Figure 5C). Likewise, NIPS significantly predicted contact heat pain sensitivity in Dataset 4: r=0.323, p=0.0003 (Figure 5C). Interestingly, this correlation was close to what the original authors of Dataset 4 observed (r=0.252) in their own study, where a specific model was built to predict contact heat pain sensitivity^17^. To test the applicability of NIPS in patient groups, we also applied it to a clinical dataset (i.e., Dataset 5). NIPS predicted pain sensitivity in both postherpetic neuralgia patients (r=0.561, p=0.024) and age- and sex-matched healthy individuals (r=0.504, p=0.024) (Figure 5C). As in the univariate analysis, we also controlled for dataset, sex, and age, and found that the revised model still predicted pain sensitivity in Datasets 1∼5 (Figure S8B&C). Note that while Datasets 1∼4 included young and middle-aged adults, Dataset 5 recruited older adults as participants (mean age>60 years). Furthermore, these datasets also included participants from different ethnicity and cultural backgrounds. Taken together, these results suggest that NIPS is a generalizable neural indicator of pain sensitivity.

To examine the effect of sample size on the model performance, we also resampled data (training sample size: 20 to 300, in steps of 5) in the pooled 3.5J condition in Datasets 1&2. Note that 5-fold cross-validation was adopted to assess the model performance. Therefore, for a sample of 100 participants, the training sample size (the abscissa in Figure 5D) is 80. When the training sample size was smaller than or equal to 70 (i.e., total sample size ≈88), NIPS’s performance was very poor, even worse than a simple mean prediction (R^2^<0), and varied dramatically. When the training sample size was around 100 (i.e., 95, 100, or 105), the R^2^ of NIPS was approximately 0.06. Interestingly, this estimate was very close to what Gim et al^17^ found with a training sample size of 99 (model performance: r=0.252, R^2^=0.064). Nevertheless, with such a sample size, NIPS’s performance still varied substantially. The performance stabilized only after the training sample size reached 120 (i.e., total N=150) and then R^2^ kept increasing with the training sample size. These findings suggest that a large sample size (>150) is needed to build a well-performing and robust pain prediction model.

While NIPS performed relatively well in our datasets, it explained only a small part of the variance of pain sensitivity. Since behavioral measures other than intensity ratings were available in Datasets 1&2, we explored whether incorporating them with fMRI responses could help build better predictive models. As a first step to build this composite model, we trained a model with behavioral measures. The behavior only model could also predict pain sensitivity: (1) Discovery Set: r=0.466, p<0.0001, R^2^=0.213; (2) Holdout Test Set: r=0.356, p<0.0001, R^2^=0.123 (Figure S9B). More importantly, utilizing both fMRI responses and behavioral measures led to a better-performing model: (1) Discovery Set: r=0.634, p<0.0001, R^2^=0.401; (2) Holdout Test Set: r=0.606, p<0.0001, R^2^=0.361 (Figure S9C). The R^2^ of the fMRI + behavior model was roughly the sum of the fMRI only model and behavior only model. While the generalizability of this composite model could not be tested due to the unavailability of behavioral measures in other datasets, our result still illustrates a promising approach for constructing more powerful composite models.

### NIPS is a distributed pattern predictive of pain reduction in healthy individuals

Having comprehensively tested NIPS’s performance, we then attempted to understand what it represents. We first used virtual lesion to examine which brain areas contributed to NIPS’s predictive power^44^. We then tested NIPS’s ability to predict the analgesic effects of placebo (Dataset 3) and neuromodulation (Dataset 6) in healthy individuals.

To explore which areas were more important for NIPS, we parcellated the brain according to the Automatic Anatomical Labeling template^45^, removed (i.e., “virtually lesioned”) each area, and refitted the predictive model without the corresponding area. Although the effect of removing areas was not uniform across the brain, none of the areas was practically indispensable for NIPS’s performance, leading to a maximum decrease of R^2^ only by 0.02 (Figure 6A; see supplemental data file 1 for complete results). However, keeping only one area while removing all others greatly reduced NIPS’s performance, causing a median drop of R^2^ by 0.29 even though the original R^2^ was only 0.24 (Figure 6B; see supplemental data file 2 for complete results). Taken together, these findings suggest that information about pain sensitivity is represented in a distributed neural system, rather than local and isolated regions^46^.

**Figure 6.**
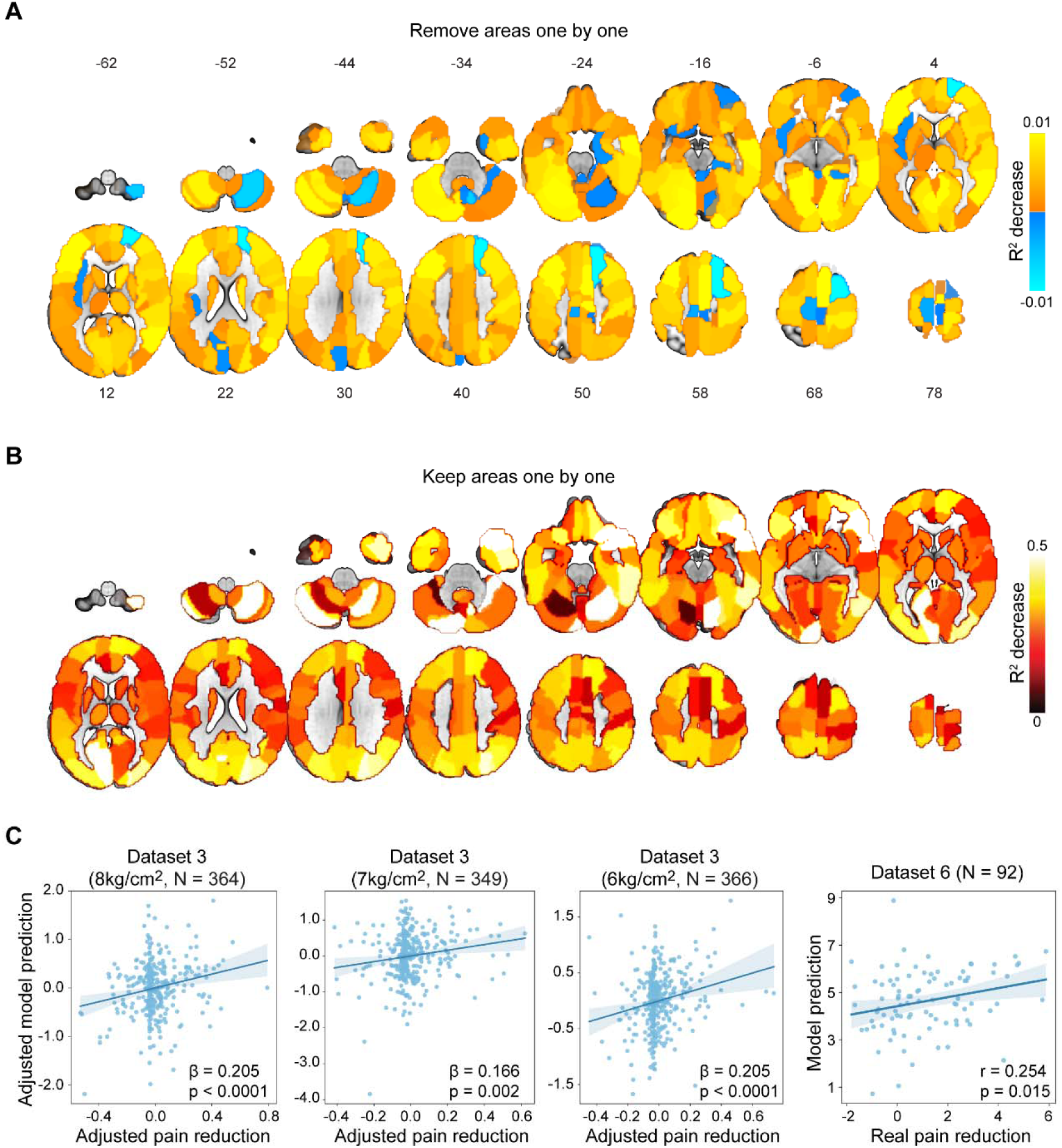
Region importance and pain reduction prediction of NIPS. **(A)** Region importance assessed by removing areas one by one in the AAL template. Removing only single brain region did not make NIPS’s performance substantially worse, suggesting that a distributed network rather than isolated areas is responsible for representing pain sensitivity. **(B)** Region importance assessed by keeping areas one by one in the AAL template. Since the R^2^ of NIPS was only about 0.24, keeping only single brain region substantially impacted NIPS’s performance, suggesting that regions work together to represent pain sensitivity. **(C)** Prediction of pain reduction by NIPS in healthy individuals. In Datasets 3&6, NIPS significantly predicted pain reduction after pain treatments, suggesting NIPS could be related to pain modulation. In Dataset 3 results, “adjusted” means that random effects of family were adjusted. β is the standardized coefficient in mixed effects models. Note that real and predicted values in Dataset 6 were not adjusted since dependent observations were not present in this datasets. Shaded regions in (C) stand for 95% confidence intervals.

To further understand what NIPS represents, we then applied NIPS to predict pain reduction in healthy individuals using Datasets 3&6. Dataset 3 was originally collected to investigate placebo analgesia^37^. We previously utilized the fMRI data in the control condition in Dataset 3 to test the generalizability of NIPS (Figure 5C). We now subtracted BOLD responses and mean pain ratings in the placebo condition from the control condition to quantify the placebo effect. NIPS significantly predicted the magnitude of placebo analgesia: (1) the 8kg/cm^2^ condition: β=0.205, p<0.0001; (2) the 7kg/cm^2^ condition: β=0.166, p=0.002; (3) the 6kg/cm^2^ condition: β=0.205, p<0.0001 (Figure 6C). Dataset 6 was collected to study the analgesic effect of transcutaneous electrical nerve stimulation (TENS) with a pretest-treatment-posttest design^38^. To quantify the analgesic effect, we subtracted BOLD responses and mean pain ratings in the posttest from those in the pretest. We also found a significant correlation between the model predictions and real pain reductions (r=0.254, p=0.015; Figure 6C). Furthermore, after accounting for the influence of baseline pain ratings (control condition in Dataset 3 and pretest condition in Dataset 5), we still found that predicted analgesic effects were significantly associated with real pain reductions: (1) the 8kg/cm^2^ condition in Dataset 3: β=0.156, p=0.0005; (2) the 7kg/cm^2^ condition in Dataset 3: β=0.101, p=0.017; (3) the 6kg/cm^2^ condition in Dataset 3: β=0.085, p=0.040; (3) Dataset 5: partial r=0.238, p=0.023. Regressing dataset, sex, and age also had no substantial effect on the ability of the multivariate model to predict pain reduction in Datasets 3 and 6 (Figure S8D). These findings converged to support the proposition that our fMRI-based pain sensitivity model predicts, at least partly, pain reduction from interventions in healthy individuals.

## Discussion

With six large and ethically and culturally diverse fMRI datasets, we systematically examined the relationship between nociceptive-evoked BOLD responses and pain sensitivity across individuals. We obtained four major findings. First, there were replicable and generalizable correlations between nociceptive-evoked BOLD responses and pain sensitivity. Second, the correlations between BOLD responses and sensory sensitivity were more evident in pain than in other modalities. Third, NIPS accurately predicted not only pain sensitivity across different pain types, but also pain relief from different pain treatments. Fourth, sample sizes had enormous impacts on assessing the relationship between BOLD responses and pain sensitivity using both univariate correlation analysis and multivariate predictive modelling. Altogether, using the largest dataset on nociceptive-evoked fMRI responses available to date, we demonstrate that BOLD responses can serve as replicable, generalizable neural indicators of pain sensitivity. These findings would facilitate interpretations of subjective pain reports and promoting more mechanistically informed investigation of pain physiology.

Understanding the neural basis of variability of pain sensitivity has been a major topic in pain neuroscience^10–14,26,47–49^. Theoretically, nociceptive-evoked BOLD responses reflect real-time neural processing of painful stimuli, thus potentially having a stronger correlation with pain sensitivity than other brain features, such as grey matter volume^47^, cortical thickness^49^, and resting-state functional connectivity^12,13^. Indeed, task fMRI data do improve the prediction of cognitive traits compared with resting-state fMRI data^50,51^. However, it is still highly controversial whether nociceptive-evoked BOLD responses can reliably encode pain sensitivity, since inconsistent findings were obtained in different groups even when the sample size (≈100) seemed large^11,17^. Leveraging six datasets with a total sample size of 1046, we found that nociceptive-evoked BOLD responses reliably correlated with pain sensitivity. This correlation was robust with respect to statistical methods used (parametric or nonparametric) and potential confounders (dataset heterogeneity, sex, and age), replicable in multiple datasets with different stimulus intensities, and generalizable across different pain types in ethnically and culturally diverse populations. In agreement with previous studies^10,15,17^, typical pain-related areas were also among the areas encoding pain sensitivity. We thus provided compelling evidence that, given sufficiently large sample sizes, pain sensitivity could be indexed from BOLD responses to pain, which is consistent in principle with previous studies showing the ability of nociceptive-evoked electrophysiological responses to encode pain sensitivity^26,48^.

We also developed NIPS to predict pain sensitivity from nociceptive-evoked BOLD responses. With multiple independent datasets, we confirmed that NIPS has good predictive performance. Cross-validation and testing in the Holdout Test Set demonstrated that NIPS explained more than 20% of the variance of pain sensitivity. Crucially, we tested NIPS’s generalizability by applying it to mechanical pain data in Dataset 3 and contact heat pain data in Dataset 4. Unsurprisingly, its performance dropped in these two external datasets, since machine learning models’ performance generally decreases in external datasets that differ greatly from the discovery set^52^. Neural mechanisms of heat pain and mechanical pain are not identical^35^, and entirely different participants were included in different datasets. Furthermore, even the rating scales were different. Datasets 3&4 adopted a 0∼1 Labeled Magnitude Scale, consisting of quasi-logarithmically spaced perceptual verbal labels rating, while the Discovery Set and Holdout Test Set employed a 0∼10 numeric rating scale, whose ratings do not seem to be logarithmically spaced^53^. All these considerations may have caused the relatively worse performance of NIPS in Datasets 3&4. Nevertheless, NIPS still exhibited significant predictive power in both datasets, demonstrating broad generalizability across different pain types. This generalizability is particularly remarkable, given the diverse backgrounds of participants in our datasets: Chinese in the Discovery and Holdout Test Sets, Americans in Dataset 3, and South Koreans in Dataset 4. NIPS thus showed extensive generalizability across ethnic groups and cultures, even though previous studies have observed ethnic and cultural differences in pain^54,55^. While participants in Datasets 1∼4 were relatively young, NIPS could also predict pain sensitivity in older individuals in Dataset 5 (mean age>60 years). More importantly, NIPS predicted pain sensitivity in PHN patients in Dataset 5, suggesting some potential clinical relevance of this model, although its performance in patients with chronic pain still requires further testing in more clinical datasets.

Given these robust and generalizable findings, why did some studies fail to find any correlation between BOLD responses and pain sensitivity? We examined two likely explanations. One is that the physical intensity of painful stimuli may be too small to evoke a clear pain sensation^56^. The power to detect correlations will be dampened if no clear pain sensation is elicited or BOLD responses are just too noisy due to insufficient physical intensity. Our results from Dataset 3 supported this explanation: fewer areas showed significant correlations with pain sensitivity as the physical intensity decreased. The other explanation is that the sample size may be simply not large enough. Traditionally, neuroimaging studies have limited sample sizes^57,58^, which is a crucial factor in the non-replicability of study findings^22,24^. Taking advantage of our large datasets, we systematically investigated the impact of sample size on the detectability of univariate correlations between nociceptive-evoked BOLD signals and pain sensitivity with resampling analysis. We found that a sample size of around 200 could only detect less than 25% of the significant voxels in univariate correlation analysis with 80% power. Similarly, sample sizes also had a substantial influence on the predictive power of NIPS. Its predictive performance improved continuously as the sample size grew, suggesting that better models can be built with even more data^59–61^. On the other hand, the performance of NIPS varied dramatically when the sample size was small. Only when the sample size reached 150 did we observe stable predictive performance. In summary, a sample size >200 seems required for univariate correlation analysis to reveal the correlation between BOLD response and pain sensitivity, and a sample size >150 may be needed to build a robust and well-performing ML model to decode pain sensitivity. This conclusion resonates with recent studies showing that a massive sample size like several hundred or even thousand is required to detect replicable brain-behavior associations^23,24,62,63^.

Previous studies have shown that painful and nonpainful stimuli activate more or less the same brain regions^27,29,32^, which was replicated in our study, suggesting that BOLD responses may not selectively reflect pain. We did find significant correlations between nonpainful stimuli-evoked BOLD responses and corresponding sensory sensitivity, which agrees with a recent study showing correlations between BOLD responses and auditory sensitivity^11^. However, we observed larger correlations between BOLD responses and pain sensitivity than between BOLD signals and tactile/auditory/visual sensitivity, even after sensitivity differences between modalities were controlled by a matching procedure. Therefore, although the correlation between BOLD responses and sensory sensitivity is not unique to pain, there are quantitatively larger correlations in pain than in other sensory modalities. In other words, the brain is more capable of representing pain sensitivity. Some recent studies have identified brain areas whose activity preferentially encodes pain^30–33^, but they have mainly focused on the relationship between fMRI responses and pain perception at the within-individual level. Our study thus advances this line of research by showing that fMRI responses differentially encode pain sensitivity at the between-individual level. Future research is in need to reveal the underlying mechanisms of differential encoding of pain sensitivity and nonpain sensitivity.

Apart from establishing the correlation between BOLD responses and pain sensitivity, the present study also deepens our understanding of neural representations of pain sensitivity. When removing brain areas one by one, we observed few to no reductions in NIPS’s predictive power. On the other hand, keeping only one area greatly impacted the predictive performance of NIPS. These findings are in agreement with real brain lesion studies showing that lesions in key pain-related areas do not disrupt pain intensity perception^64,65^. Pain sensitivity thus is represented in a distributed neural system, rather than local and isolated regions^46^.

One particularly interesting finding was that NIPS could predict individual differences in pain reduction from different interventions in healthy participants. Since NIPS also predicted individual differences in pain sensitivity, these two types of individual differences may be related, which is supported by previous studies showing significant associations between pain sensitivity and conditioned pain modulation^66,67^. Neurobiologically, the perception of pain reflects both the ascending nociceptive input and descending pain modulatory system^68^. Our findings thus could mean that pain sensitivity may be partially determined by individuals’ ability to modulate pain. This statement is in line with the finding that NIPS explained more variance of pain sensitivity than the variance of analgesic effects in the same independent dataset not involved in the development of NIPS (i.e., Dataset 3). Alternatively, NIPS’s ability to predict pain reduction could also reflect partial overlapping of the pain modulation system in the brain and pain processing circuitry such as the rACC^69^. Note that, due to lack of clinical data, NIPS’s prediction of pain reduction in healthy individuals cannot be interpreted as NIPS can predict analgesic effects in patients with pain.

Our study has some limitations. First, we only investigated the influence of stimulus intensity and sample size on the detectability of the correlation between BOLD responses and pain sensitivity due to the nature of available datasets. Other contributing factors like sample personal characteristics and study design need to be tested in future studies. Second, while NIPS has reasonably good performance in predicting pain sensitivity, its predictive power can be further improved. As suggested by our resampling analysis, collecting even more data has the potential to boost NIPS’s predictive performance^70^. The improved performance of our composite model integrating fMRI and behavioral measures also suggest that incorporating multimodal data may be helpful to capture more variability of pain sensitivity^61^. Since fMRI alone may not be sufficient to explain the majority of pain sensitivity, data from other imaging modalities are most likely indispensable to build a model powerful enough to be applicable in real life. Finally, the clinical implications of our study are limited, since participants in our study were mainly healthy individuals. Clinical populations are typically more variable in terms of age, socioeconomic background, education, diagnosis, and symptoms. While our datasets, especially the PHN dataset (i.e., Dataset 5), exhibit some variability in age, ethnicity, cultural background, and pain diagnosis, our findings are mainly based on rather homogenous samples, and thus still need to be replicated in more heterogenous clinical populations. Although we demonstrated NISP’s ability to predict pain sensitivity in PHN patients, it remains to be tested more extensively in other chronic pain patients, and it is of particular interest to test whether NIPS can predict pain relief in patient groups.

## Methods

### Datasets overview

We utilized six fMRI datasets that had been published recently to study other research questions (Figure 1A). We collected Datasets 1, 2, 5, and 6 using E-Prime (ver. 2.0, PST Inc.) on our own in China. Datasets 3 and 4 were collected using Matlab (ver. 2018b) in the US and South Korea, respectively, and kindly shared by the original authors^17,37^. In Datasets 1 and 2, participants received laser pain, electro-tactile, auditory, and visual stimuli^14^. Dataset 3 was originally collected to investigate the neural underpinnings of placebo analgesia^37^. The researchers delivered both contact heat pain and mechanical pain stimuli to participants, but used contact heat pain stimuli to induce placebo analgesia by conditioning. To avoid potential influence of the placebo induction on the neural processing of contact heat pain, we analyzed data only for the mechanical pain stimuli in the placebo and control conditions, and discarded data with contact heat pain stimuli. Instead, we used data with contact heat pain stimuli from Dataset 4^17^, where no additional interventions like placebo were introduced. Dataset 5 delivered laser heat pain stimuli to postherpetic neuralgia (PHN) patients and healthy controls^36^. Dataset 6 also used contact heat pain stimuli, and was from a study aiming to reveal the spino-cortical mechanisms of TENS-induced analgesia^38^. In Datasets 1∼5, the physical intensity of painful stimuli was fixed. In Dataset 6, however, the physical intensity was adjusted individually for each participant. Therefore, we did not include Dataset 6 in most analyses, but only applied NIPS to it to test whether NIPS could predict pain reduction. We summarized basic information for these six datasets in Table S1.

### Participants

Dataset 1 included 212 participants. One participant did not provide the necessary demographic information. For the rest 211 participants, there were 135 females, aged 21.5 ± 4.2 years (Mean ± SD). Dataset 2 had 187 participants. There was also one participant who did not provide the demographic information. For the rest 186 participants, there were 103 females, aged 21.0 ± 3.3 years. Note that Datasets 1&2 were collected simultaneously, not separately, in a single project. Treating them as two datasets reflects the fact that participants in these datasets had different pain sensitivity as assessed in a calibration phase and received painful stimuli of different intensities to avoid delivering pain stimuli that could be potentially intolerable for participants with high pain sensitivity. In other words, participants were recruited on a rolling basis and assigned to either Dataset 1 or Dataset 2 based on their pain sensitivity, which was assessed during the calibration phase. Dataset 3 included 395 participants (231 females, age: 35.4 ± 2.6). Note that some participants in this dataset had missing behavioral ratings in some conditions. Considering the lack of sufficient trials per condition, we discarded data from these participants with missing values, which was the reason why the sample sizes in the main figures were 369, 358, and 371 for the three intensity conditions respectively. Dataset 4 had 124 participants (61 females, age: 22.2 ± 2.7). Dataset 5 included 16 right-handed patients (5 males; age [mean ± SD] = 65.75 ± 6.99 years) suffering from one condition of chronic pain called postherpetic neuralgia (PHN) and 20 age- and gender-matched right-handed healthy controls (8 males; age: 61.55 ± 8.21 years). Dataset 6 included 92 participants (50 females, age: 21.9 ± 3.2). All participants except for Dataset 5 were healthy and free of pain, neurological, or psychiatric disorders. Patients in Dataset 5 fulfilled the International Association for the Study of Pain (IASP) criteria for PHN and were diagnosed by experienced clinicians based on clinical symptoms (including medical history, shingles history, pain severity, and pain types). None of participants in Dataset 5 had a past or current diagnosis of any psychiatric or major neurological illness. All participants gave written informed consent prior to the experiments. Local ethics committees approved the experimental procedures for the original studies (Datasets 1, 2, 5, and 6: Ethics Committee of the Institute of Psychology at the Chinese Academy of Science; Dataset 3: Institutional Review Board of the University of Colorado Boulder; Dataset 4: Sungkyunkwan University Institutional Review Board).

### Sensory stimulation

Three different types of painful stimuli were used in these datasets: (1) laser heat pain: Datasets 1, 2, and 5; (2) mechanical pain: Dataset 3; (3) contact heat pain: Datasets 4 and 6. Datasets 1 and 2 also delivered three kinds of nonpainful stimuli: (1) electro-tactile, (2) auditory, and (3) visual stimuli.

In Datasets 1 and 2, laser heat pain stimuli were transient radiant heat pulses (wavelength: 1.34μm; pulse duration: 4ms) generated by an infrared neodymium yttrium aluminum perovskite (Nd: YAP) laser (Electronical Engineering, Italy). The laser beam was transmitted by an optic fiber, and its diameter was set at approximately 7mm. Laser pulses were delivered to a pre-defined square (5×5cm^2^) on the left-hand dorsum. After each stimulus, the laser beam was displaced by approximately 1cm in a random direction to avoid nociceptor fatigue or sensitization. Two stimulus energies (3.0J and 3.5J) were used in Dataset 1, and two other stimulus energies (3.5J and 4.0J) were used in Dataset 2. A stimulus intensity of 3.0J was adopted in Dataset 1 because participants in this dataset had higher pain sensitivity and nociceptive laser stimuli of 4.0J could be unbearable for them. In Dataset 3, mechanical pain stimuli were delivered to the fingers of the left hand for 10s with an MRI-compatible pressure pain device. There were three levels of physical intensity, 6 kg/cm^2^, 7 kg/cm^2^, and 8kg/cm^2^. Note that the original authors sharing Dataset did not specify the location of stimulation (e.g., nails) ^37^. We thus cannot provide more details regarding the location of mechanical stimulation. In Datasets 4, contact heat pain stimuli were administered to the volar surface of the left forearm with a 16×16mm^2^ ATS thermode of a Pathway system (Medoc Ltd, Ramat Yishai, Israel). The target temperature of the stimuli was 47.5°C with the baseline temperature being 32°C. The stimulation lasted for 12s (ramp-up: 2.5s; plateau: 7s; ramp-down: 2.5s). In Dataset 5, laser heat pain stimuli were also transient radiant heat pulses. Stimulus parameters were similar to those in Datasets 1 and 2, except that the laser energy was only set to 3.5J. In Dataset 6, contact heat pain stimuli were delivered to the C5∼C6 dermatomes on both forearms with a 573mm^2^ (diameter of 27mm) CHEPS thermode using a Pathway system (Medoc Ltd, Ramat Yishai, Israel). The heat stimulation ramped up from a baseline of 32°C to a temperature for 3s that led to a perceived intensity rating of 7 on a 0∼10 rating scale. In other words, the target temperature was tailored individually for each participant to ensure a constant perceived intensity.

In Datasets 1 and 2, nonpainful stimuli were also delivered. Non-nociceptive tactile stimuli were constant current square-wave electrical pulses (duration: 1ms; model DS7A, Digitimer, UK) delivered through a pair of skin electrodes (1cm interelectrode distance) placed on the left wrist, over the superficial radial nerve. The same two stimulus intensities (2.0mA and 4.0mA) were used in both datasets. Auditory stimuli were brief pure tones (frequency: 800Hz; duration: 50ms; 5ms rise and fall time) delivered through a headphone. The same two stimulus intensities (76dB SPL and 88dB SPL) were used for all participants in both datasets. Visual stimuli were brief flashes of a grey round disk in a black background (duration: 100ms) on a computer screen. The stimulus intensities were adjusted using the greyscale of the round disk, corresponding to RGB values of (100, 100, 100) and (200, 200, 200), respectively, for all participants in both datasets. Stimulus intensities of tactile, auditory, and visual stimuli were determined based on a pilot behavioral experiment to ensure that the perceived ratings of low and high intensity stimuli were approximately 4 and 7 out of 10, respectively.

### Rating scale

Two ratings scales were used for different datasets: 0∼10 Numeric Rating Scale and 0∼1 Labeled Magnitude Scale. In Datasets 1, 2, 5, and 6, the 0∼10 Numeric Rating Scale was used, where 0 meant ‘‘no sensation’’, and 10 ‘‘the strongest sensation imaginable’’. This is a common rating scale in pain research, but evidence has suggested that it is not a ratio scale, namely, a rating of 4 may not be exactly twice as painful as a rating of 2^53^. In Datasets 3 and 4, the 0∼1 Labeled Magnitude Scale was adopted. In Dataset 3, 0 on the scale meant “no pain”, 0.014 “barely detectable”, 0.061 “weak pain”, 0.172 “moderate pain”, 0.354 “strong pain”, and 1 “most intense sensation imaginable” ^37^. In Dataset 4, 0 meant “no sensation”, 0.061 “weak”, 0.172 “moderate”, 0.3454 “strong”, 0.533 “very strong”, and 1 “strongest imaginable”^17^. The Labeled Magnitude Scale was not used as often as the Numeric Rating Scale in pain research, but seems to be a ratio scale^71^. All participants were asked to rate the intensity of pain according their authentic sensory experience without any other considerations. Note that we simply analyzed preexisting datasets. As a result, we had no control over which rating scale to use. Additionally, due to differences in the perceptual meaning of ratings, we could not multiply the 0∼1 scale by 10 to convert it to a 0∼10 scale.

### Study design

The original authors reported their study designs in great detail^14,17,36–38^. Since these papers were all openly accessible, here we only briefly described the study design of each dataset (Figure 1B).

In Datasets 1 and 2, participants first underwent a quantitative sensory testing and filled out a series of questionnaires (see Zhang et al.^72^ for details). The quantitative sensory testing assessed laser heat pain threshold, and cold pain threshold and tolerance. Questionnaires included the Chinese versions of Fear of Pain Questionnaire (FPQ), Pain Anxiety Symptoms Scale-20 (PASS), Pain Catastrophizing Scale (PCS), Pain Vigilance and Awareness Questionnaire (PAVQ), Beck Depression Inventory (BDI), Behavioral Activation and Behavioral Inhibition Scales (BAS/BIS), Revised Life Orientation Test (LOT-R), Pittsburgh Sleep Quality Index (PSQI), Interpersonal Reactivity Index (IRIC), Big Five Personality Test, and State-Trait Anxiety Inventory (STAI). Then participants received 80 transient stimuli of four different sensory modalities (nociceptive laser, non-nociceptive tactile, auditory, and visual) divided into two runs, and then rated their perceived intensity with a rating scale ranging from 0 (‘‘no sensation’’) to 10 (‘‘the strongest sensation imaginable [in each stimulus modality]’’). For each sensory modality, two stimulus intensities (i.e., high and low) were delivered. In other words, each participant underwent eight experimental conditions (4 modalities×2 intensities), each condition with 10 trials. The only difference in the study design of Datasets 1 and 2 was the physical intensity of laser heat pain stimuli (detailed in **Sensory stimulation**).

Dataset 3 adopted a within-subjects design to examine the neural mechanisms of placebo analgesia. The study consisted of a verbal suggestion phase, a conditioning phase, and a test phase. Verbal suggestions and conditioning were used to induce placebo effects. In the verbal suggestion phase, two identical creams were introduced to participants, one being a placebo cream that was claimed to have a pain-relieving effect, the other being a control cream claimed to have no analgesic effects. These two creams were applied to two different fingers of the left hand. Then participants went through the conditioning phase, where they experienced the analgesic effect of the placebo cream through social observational learning and classical conditioning. Importantly, in this conditioning phase, only contact heat pain stimuli were delivered. Finally, in the test phase, participants received both contact heat pain and mechanical pain stimuli in the fingers with the placebo and control creams. The test phase delivered a total of 32 stimuli in four runs, with four thermal and four mechanical stimuli in a random order per run. After each stimulus, participants rated the intensity of the stimulus with a 0∼1 Labeled Magnitude Scale, consisting of quasi logarithmically spaced perceptual verbal labels rating scale, where 0 means “no pain”, 0.014 “barely detectable”, 0.061 “weak pain”, 0.172 “moderate pain”, 0.354 “strong pain”, and 1 “most intense sensation imaginable”.

In Dataset 4, participants received contact heat pain stimuli of six intensity levels in their left forearm. The entire study consisted of eight pain task runs with 12 trials each run and 16 trials per temperature. After each stimulus, participants rated the perceived intensity of the stimuli with a 0∼1 Labeled Magnitude Scale. Note that the original authors only shared data for the highest temperature condition (i.e., 47.5°C). We thus could only analyze data for this condition.

In Dataset 5, participants received 20 laser heat pain stimuli and reported their perceived intensity and unpleasantness ratings with a 0∼10 Visual Analog Scale (VAS), with 0 indicating “no sensation” and 10 indicating “unbearable sensation”. We only analyzed intensity ratings to focus on the sensory aspect of pain.

Different from Datasets 1∼5, Dataset 6 adopted a between-subjects design to investigate the cortico-spinal mechanisms of TENS-induced analgesia. Participants were randomly assigned to three groups: conventional TENS (c-TENS), acupuncture-like TENS (a-TENS), and sham TENS. Each participant experienced 60 contact heat pain stimuli (pre-treatment session: 15 stimuli in the left arm and 15 stimuli in the right arm; post-treatment session: 15 stimuli in the left arm and 15 stimuli in the right arm) in two runs and then reported their perceived intensity and unpleasantness in a 0∼10 ratings scale (0: “no feeling/unpleasantness”; 10: “unbearable pain/extreme unpleasantness”). Note that we only used data from the 15 stimuli delivered to the left arm, since painful stimuli were only delivered to the left hand in other datasets. We pooled data from all three groups. The purpose was twofold. First, we could have a large sample size and thus a larger statistical power. Second, our goal was to test whether NIPS could predict the amount of pain reduction after the treatment compared with the pre-treatment baseline, not the analgesic effect of verum TENS compared with sham TENS. Indeed, pain sensitivity could also change after the sham TENS treatment due to factors like natural history, regression to the mean, and so on. A model that predicts pain reduction should thus also predict sensitivity change in the sham TENS condition.

### MRI acquisition

Details of the MRI acquisition parameters can be found in Table S2&S3.

### Image preprocessing

We analyzed preprocessed data from the original studies^14,17,36–38^. Detailed preprocessing descriptions can be found in these papers. For Datasets 1&2, fMRI data were preprocessed using Statistical Parametric Mapping 12 (SPM12) (Wellcome Trust Center for Neuroimaging, London). The preprocessing included: (1) removing the first three volumes in each run; (2) slice-time correction using the second slice and realignment to the mean slice; (3) co-registration; (4) normalizing to the Montreal Neurological Institute (MNI) space (resampling voxel size = 3×3×3mm^3^); (5) regressing out five principal components of the white matter (WM) and cerebrospinal fluid (CSF) signals, and the six motion parameters; (6) smoothing with a 6mm full-width at half maximum (FWHM) Gaussian kernel.

For Dataset 3, images were preprocessed with fMRIprep ver. 20.2.3^73^. Briefly, the preprocessing included: (1) motion-correction with MCFLIRT (FSL 5.0.9); (2) co-registering to the T1 reference using the boundary-based registration method (BBR) (FreeSurfer); (3) normalizing to the standard space; (4) smoothing with a 6mm FWHM Gaussian kernel.

For Dataset 4, preprocessing was performed with SPM12 and FSL, including (1) removing the first 18 volumes for each run; (2) realignment; (3) co-registration; (4) normalizing to the MNI space (2×2×2 mm³); (5) smoothing with a 5mm FWHM Gaussian kernel; (6) reducing motion-related artifacts with the Independent Component Analysis-based strategy for Automatic Removal Of Motion Artifacts; (7) removing data where the mean frame displacement (FD) of a run > 0.2mm, or the FD of any volume > 5mm.

For Dataset 5, preprocessing was conducted with FSL, including (1) motion correction using MCFLIRT; (2) removal of non-brain structures using Brain Extraction Tool; (3) spatial smoothing using a Gaussian kernel with a 5mm FWHM, and high-pass temporal filtering (cut off: 100 s); (4) Independent component analysis (ICA)-based denoising was performed for each individual fMRI data to remove the artifacts, including head motion, white matter and cerebrospinal fluid noise, high-frequency noise, slice dropouts, gradient instability, EPI ghosting, and field inhomogeneities.

For Dataset 6, preprocessing was also conducted using FSL, with the following steps: (1) physiological noise (i.e., respiratory and cardiac noise) regression with the physiological noise model toolbox; (2) distortion correction with the TOPUP tool; (3) slice timing correction with the *slicetimer* function; (4) motion-correction with MCFLIRT; (5) brain extraction with the BET; (6) smoothing with a 5mm FWHM Gaussian kernel; (7) high-pass filtering (cutoff: 100s); (8) co-registration with the BBR; (9) normalizing to the MNI space (2×2×2 mm³) with FNIRT.

### fMRI data analysis

#### First level analysis

We mainly reanalyzed first level contrast images from the original studies^14,17,36–38^. In Datasets 1&2, regressors in the first level analysis included eight conditions (4 modalities×2 intensities) convolved with the canonical hemodynamic response function, its temporal derivatives, and six head motion estimates. Moreover, we high-pass filtered images with a cutoff period of 128s and accounted for temporal autocorrelations using the first-order autoregressive model (AR(1)). In Dataset 3, regressors in the first level analysis included all experimental conditions convolved with the canonical hemodynamic response function, 24 motion parameters, and mean CSF signal. A high-pass filter of 180s was also applied. In Dataset 4, a single trial approach was utilized. Regressors included every single pain trial convolved with the canonical hemodynamic response function, five principal components of WM and CSF signal, and a linear trend. Signals were also filtered with an 180s high-pass filter. Trials with a variance inflation factor > 3 were subsequently removed. Single trial beta maps were finally averaged to derive the beta map for painful stimulation. In Dataset 5, regressors included the onset of laser stimuli and rating period convolved with a gamma hemodynamic response function and their temporal derivatives. In Dataset 6, regressors included painful stimulation and rating period convolved with the canonical hemodynamic response function and its temporal derivatives.

#### Second level analysis

In Datasets 1&2, to examine the relationship between nociceptive-evoked BOLD responses and pain sensitivity, we correlated first level beta maps in each pain condition with average pain ratings. The significance threshold was set at p(FDR)=0.05 (two-tailed). To test the robustness of results, we computed both Pearson’s r and Spearman’s rho in the correlational analyses. In addition to whole brain analysis, we also conducted an ROI analysis. Specifically, we extracted mean signals from classical pain-related areas^15,16^, namely, the S1, S2, ACC, insula, and thalamus using anatomically defined masks defined with the Harvard-Oxford Atlas distributed with FSL^74^. Note that the Harvard-Oxford Atlas is a probabilistic atlas and we thresholded probability maps for each region at 50%^75^.

Although Datasets 1&2 were assumed to differ only in the pain sensitivity and laser stimulation parameters, there might still be some unintended systematic differences between them. To control the potential influence of this dataset differences, sex, and age, we conducted a revised partial correlation analysis while controlling for dataset, sex, and age. Specifically, we first standardized BOLD contrast estimates voxel-wise and behavioral measures for each dataset separately, then regressed out dataset (dummy coded as 0 and 1 for Datasets 1 and 2), sex (dummy coded as 0 and 1), and age effects from BOLD contrast estimates and mean pain ratings, and finally correlated the residuals of BOLD contrast estimates and mean pain ratings. With this dataset-wise standardization procedure, we accounted for not only the difference in the mean of ratings and BOLD responses but also the difference in the range of these variables. Note that this revised partial correlation analysis is equivalent to standardize variables dataset-wise and run multiple regression with the standardized variables. Datasets 1&2 had two missing values in sex and age, which were imputed with the median value.

To test the pain selectivity of the correlation between BOLD responses and sensory sensitivity, we also performed the same correlational analysis for touch, audition, and vision in Datasets 1&2. To directly test the significance of correlation differences, we compared the correlation coefficients between modalities^42^. The statistic for this comparison is defined as

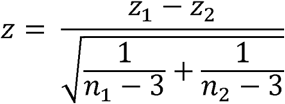

where *z_1_*and *z_2_* are the Fisher Z-transformed correlation coefficients, and *n_1_* and *n_2_* are the sample sizes.

Some participants in Dataset 3 were twins from the same family. To account for this observation dependence introduced by the family factor, we ran mixed effects models with random intercepts of family. Specifically, we adopted the following model (in Wilkinson notation): *Pain ∼ BOLD + (1|family_ID)*. To control for the potential influence of sex and age, we also run the following additional mixed effects model: *Pain ∼ BOLD + sex + age (1|family_ID)*. The significance threshold was set at p(FDR)=0.05 (two-tailed). As in Datasets 1&2, we also conducted an ROI analysis of the S1, S2, ACC, insula, and thalamus using anatomically defined masks defined with the Harvard-Oxford Atlas distributed with FSL.

Note that all statistical tests in the article are two-sided unless stated otherwise.

### Sensitivity matching procedure

Although the physical intensity of sensory stimuli was fixed in Datasets 1&2, perceived intensity ratings (i.e., sensory sensitivity) were significantly different among different modalities. Distinct correlational results in pain and nonpain modalities could thus be potentially attributed to intensity rating differences. Furthermore, perceived intensity ratings are highly correlated with stimulus salience^33^. As a result, differences in perceptual ratings may result in incompatible salience levels between modalities. To control for this possible confounding factor of rating differences, we adopted a rating matching procedure to equalize intensity ratings between pain and nonpain modalities. To rule out the influence of individual differences, we matched pain intensity ratings and tactile/auditory/visual intensity ratings in a within-individual manner. Specifically, we manually screened participants whose pain intensity ratings in the 3.5J condition and tactile/auditory/visual intensity ratings in the high intensity condition were not significantly different, namely the 95% confidence interval of the rating differences contained 0. With this matching criterion, 192 participants were selected for the pain-touch comparison, 171 for the pain-audition comparison, and 147 for the pain-vision comparison (Figure 5A∼C).

### Resampling procedure

We used a resampling method to examine the influence of sample sizes on the probability of detecting significant correlations between BOLD responses and pain sensitivity. In the pooled 3.5J data from Datasets 1&2 (N=399), we generated bootstrapped subsamples from the whole dataset, that is, randomly selected data samples with replacement. The sample size of these subsamples ranged from 100 to 400 (in steps of 10), and, for each sample size, the resampling procedure was repeated 100 times. In each subsample, we correlated BOLD responses with pain sensitivity and thresholded the correlation maps with p(FDR) = 0.05. Since we generated 100 subsamples for each sample size, the probability of a voxel being significant under a certain sample size can be approximated as the number of times that voxel reached significance. We then calculated the sample size needed for each voxel to pass the p(FDR) = 0.05 threshold with a probability of 0.8^39^.

To supplement this whole brain analysis, we also conducted the resampling in an ROI analysis manner. Specifically, we defined anatomical masks for the S1, S2, ACC, insula, and thalamus using the Harvard-Oxford Atlas^74^ distributed with FSL. Note that the probability maps for each region were thresholded at 50%^75^. Subsequently, we generated 1000 bootstrapped subsamples with sample sizes from 10 to 400 (in steps of 10). In this ROI analysis, we set the significance to α = 0.01 or α = 0.001, and then made line charts to fully show the relationship between sample sizes and the probability of significance for each ROI.

### Machine learning

#### Model development

We split randomly the pooled 3.5J data in Datasets 1&2 into a Discovery Set (N = 199) and a Holdout Test Set (N = 200). A LASSO-PCR model was trained to predict pain sensitivity using first level t maps in the Discovery Set with Scikit-learn (ver 1.3.0; https://scikit-learn.org/stable/index.html). The LASSO is a regularization method that shrinks the linear regression coefficient estimates toward zero^76^. Mathematically, LASSO regression minimizes the following loss function:

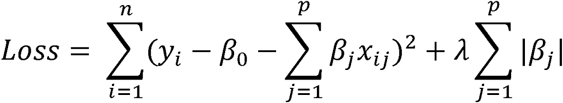

where *y_i_* is the predicted variable of participant *i*, *β*_0_ and *β_j_* are the linear regression coefficient estimates, *x_ij_* is the *j*-th predictor for participant *i*, λ is the tuning parameter, *n* is the number of participants, and *p* is the number of predictors.

T maps were first standardized across participants and then submitted to principal component analysis to reduce the feature dimensions. We retained all principal components and used them as features in the LASSO regression model (Figure 6A). Five-fold cross-validation was used to evaluate model performance. The predictive performance of the final model was assessed using two metrics: (1) Pearson’s correlation coefficient between the predicted and real pain sensitivity, and (2) R^2^, a metric representing the proportion of variance explained by the model. R^2^ is defined as:

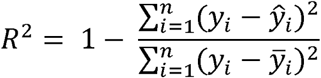

where *y_i_* is the real value for *i*-th data point, *ŷ_i_* is the predicted value, and *y̅_i_* is the mean of all real values. Note that R^2^ is not identical to the square of Pearson’s correlation, and can be negative if model performance is worse than a simple mean prediction, that is, using *y̅_i_* as predictions for all data points.

To determine the optimal tuning parameter λ, we selected 100 λ values log-uniformly distributed in the [10^−2^, 10^2^] interval and tuned λ according to the cross-validated mean squared errors. The optimal λ was 0.236.

To examine the influence of dataset heterogeneity in the pooled 3.5J data in Datasets 1&2, we regressed out dataset, sex, and age effects from first level t maps and mean pain ratings (i.e., pain sensitivity), and built a LASSO-PCR model. Note that to avoid data leakage issues, we did not standardize t maps and mean pain ratings within dataset separately, but only regress out the effect of dataset (dummy coded as 0 and 1 for Datasets 1 and 2), sex, and age. Five-fold cross-validation was also used to evaluate model performance and tune the parameter λ from 100 values log-uniformly distributed in [10^−2^, 10^2^]. The optimal λ was 0.01.

#### Model testing

After our model (i.e., NIPS) was developed, we tested its performance in the Holdout Test Set. NIPS was developed with laser heat pain data in Datasets 1&2. To test its generalizability, we then applied the model to mechanical pain data in Dataset 3 and contact heat pain data in Dataset 4. Note that the original authors of Datasets 3&4 shared beta maps for pain, not t maps. NIPS’s performance in these two datasets may thus be slightly affected due to scaling issues in beta maps. To test NIPS’s ability to predict pain sensitivity in chronic pain patients, we applied it to Dataset 5, which including PHN patients.

To examine the influence of sample size on NIPS’s performance, we refitted the model with different training sample sizes with the “*learning_curve*” function in Scikit-learn. The *learning_curve* function generates a curve showing how the performance of the model changes as the training sample size grows. We first randomly shuffled the pooled 3.5J data in Datasets 1&2, and then inputted the model and the shuffled dataset to the *learning_curve* function. We considered training sample sizes from 20 to 300 (in steps of 5). This whole process was repeated 100 times to also estimate the variability of the relationship between training sample size and model performance.

#### Model interpretation

To understand each brain area’s contribution to our model, we employed the virtual lesion approach^44^. Specifically, we parcellated NIPS’s weight map according to the Automatic Anatomical Labeling (AAL) template^45^ and removed areas one by one or kept areas one by one to examine each area’s contribution to the model performance. To assess the potential association of NIPS with pain modulation, we also applied NIPS to Datasets 3&6 to predict their analgesic effects from different pain treatments in healthy individuals. To address observation dependence in Dataset 3, we adopted the following model to evaluate the relationship between real and predicted pain reduction: *real_value ∼ predicted_value + (1|family_ID)*. To control for the potential influence of sex and age, we also run the following additional mixed effects model: *real_value ∼ predicted_value + sex + age (1|family_ID)*.

#### Ensemble model combining fMRI and behavior measures

To assess whether fMRI data together with behavioral data contribute to a composite model that better explains variance of pain sensitivity, we trained an ensemble model that included both fMRI and behavior measures. The behavior measures encompassed quantitative sensory testing and questionnaires (see supplemental data file 3 for details). As a first step to build this ensemble model, we trained a behavior only model apart from the fMRI only NIPS model. Since the number of features in the behavior only model was limited, we omitted the PCA step, and only standardized behavioral features and then input them to the LASSO algorithm. The optimal λ was selected from 100 values in [10^−2^, 10^2^] by five-fold cross-validation, and turned out to be 0.196. The fMRI + behavior ensemble model was a stacking regression model^77^. Specifically, the predictions from the fMRI model (LASSO-PCR) and behavior model (LASSO) were used as features in a final multiple regression model. This stacking model had two tuning parameters λ_1_ and λ_2_ for the embedded fMRI model and behavior model respectively. Since all combinations of λ_1_ and λ_2_ need to be explored in a grid search, training stacking models is computationally intensive. We thus only selected 10 values log-uniformly distributed in [10^−2^, 10^2^] for both λ_1_ and λ_2_. Five-fold cross-validation led to the following optimal parameters: λ_1_ = λ_2_ = 0.215. Note that while this stacking model had better performance (Figure S9), behavioral measures were not available in other datasets except for Datasets 1∼2, resulting in our inability to test the generalizability of the fMRI + behavior model.

## Data and code availability

The data and code for all results will be made available on the Open Science Framework (OSF) upon publication. Source data and code for the development and test of NIPS are available on OSF (https://osf.io/y2n34/). Run each cell in “NIPS development and test.ipynb” to reproduce how NIPS was developed and tested.

## Acknowledgment

We thank Yu-Xin Chen, Xin Hou, Xiao-Yun Li, Guang-Yue Tian, Wen-Xin Su, Fei-Xue Wang, Jia-Hui Zhong, and Yu-Pu Zhu for their assistance in data collection. We also thank colleagues collecting Datasets 3&4 for kindly sharing their data. This work was supported by the National Key Research and Development Program of China (2023YFC2508702), National Natural Science Foundation of China (32071061), and Beijing Natural Science Foundation (JQ22018).

## Author Contributions

Conceptualization: LBZ, LH; Methodology: LBZ, LH; Investigation: HJZ, LH, ZXW, LBZ; Formal analysis: LBZ, LH; Visualization: LBZ, LH; Writing – original draft: LBZ; Writing – review & editing: LBZ, XJL, HJZ, ZXW, YZK, YHT, GDI, LH; Funding acquisition: LH; Project administration: LH; Supervision: LH.

