## Supplemental figs and tables for "A replicable and generalizable neuroimaging-based indicator of pain sensitivity across individuals"

**Supplemental figures**

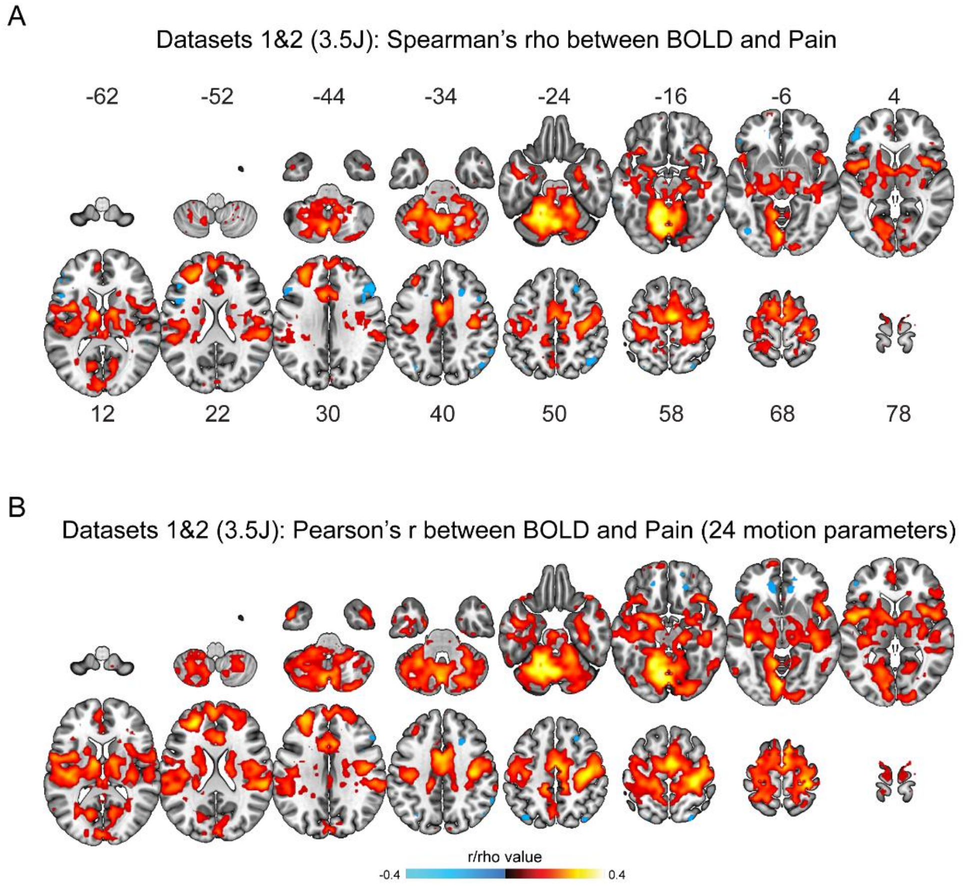


**Figure S1. Robustness of the correlation between BOLD responses and pain sensitivity in Datasets 1&2. (A)** Nonparametric Spearman correlation between nociceptive-evoked BOLD responses and pain sensitivity in Datasets 1&2. Nonparametric correlation revealed similar correlational patterns as parametric correlation. **(B)** Correlational results when head motion was controlled with 24 parameters in Datasets 1&2. Strict control of head motion did not substantially change the correlational patterns between BOLD responses and pain sensitivity.

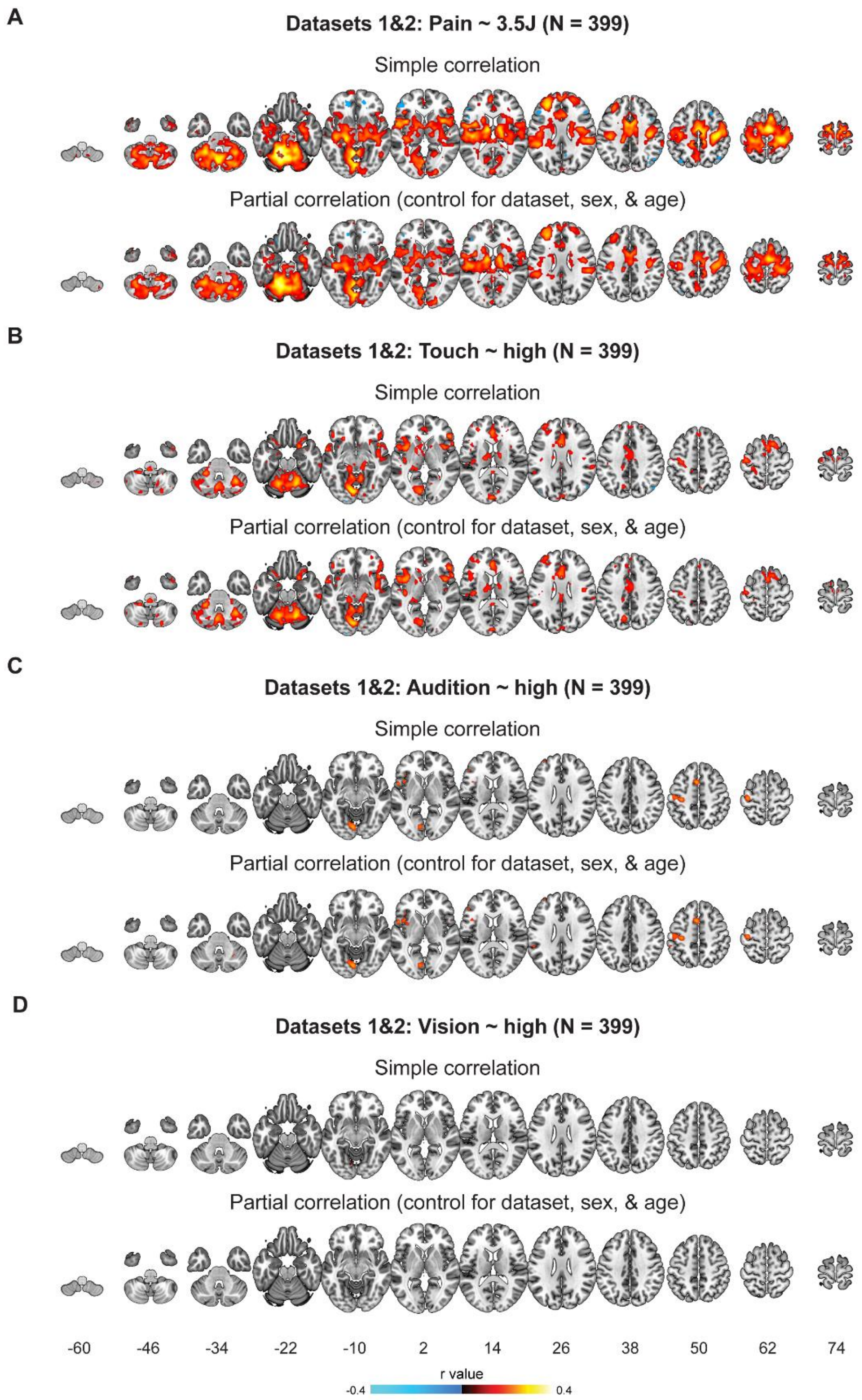


**Figure S2. Comparisons of results between simple correlations and partial correlations for different sensory modalities in Datasets 1&2.** Simple correlation results were identical to those reported in Figures 2 and 4. Partial correlation results were controlled for dataset (dummy coded as 0 and 1), sex (dummy coded as 0 and 1), and age. Controlling for dataset, sex, and age slightly decreased the magnitude of correlations between fMRI response and sensory sensitivities, but the general pattern remained the same.

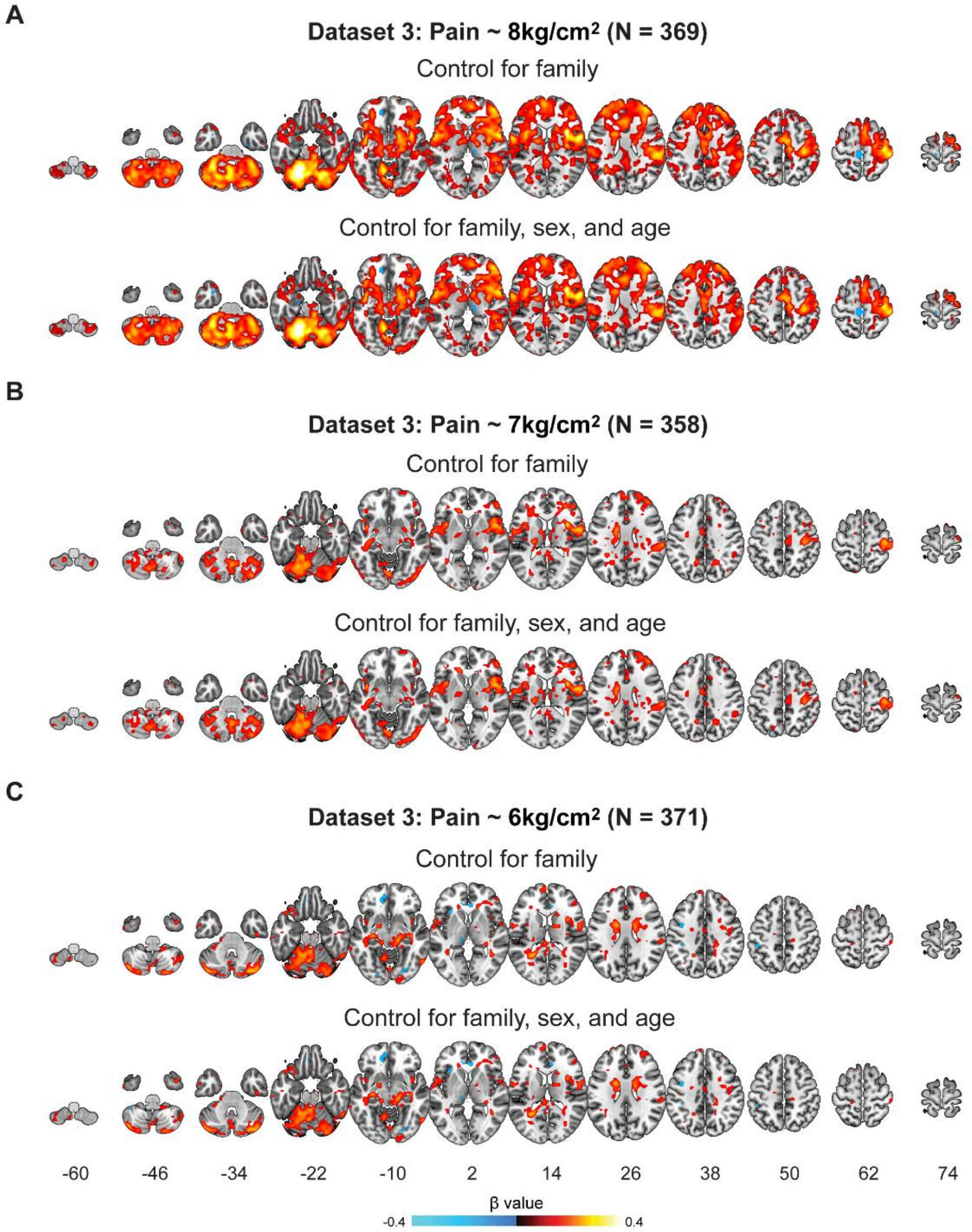


**Figure S3. Results before and after controlling for sex and age in Dataset 3.** Further control over sex and age had no substantial effect on the associations between fMRI responses and mechanical pain sensitivity.

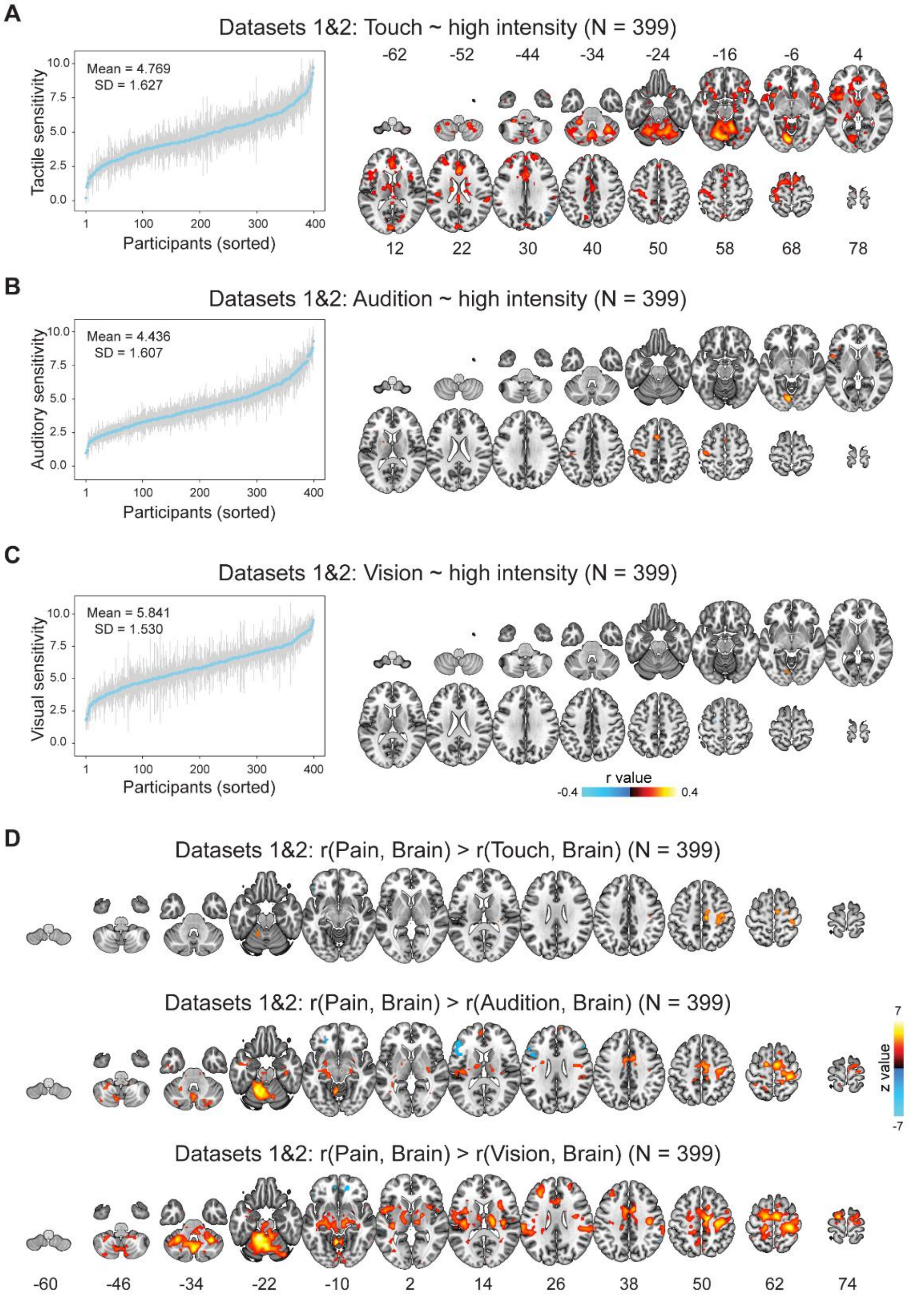


**Figure S4. Correlations between BOLD responses and sensory sensitivity in nonpain modalities. (A)~(C)** BOLD responses-sensory sensitivity correlation in the high intensity condition in Datasets 1&2 for touch (A), audition (B), and vision (C). Inter-individual variability of tactile sensitivity correlated with BOLD responses in many areas, while only a few areas showed correlation with auditory and visual sensitivity. **(D)** Comparisons of BOLD-sensitivity correlation between pain and nonpain modalities. Direct comparisons of correlation coefficients showed larger correlations for pain than for touch, audition, and vision. Error bars in (A)~(C) stand for standard deviation (SD) of intensity ratings for each participant.

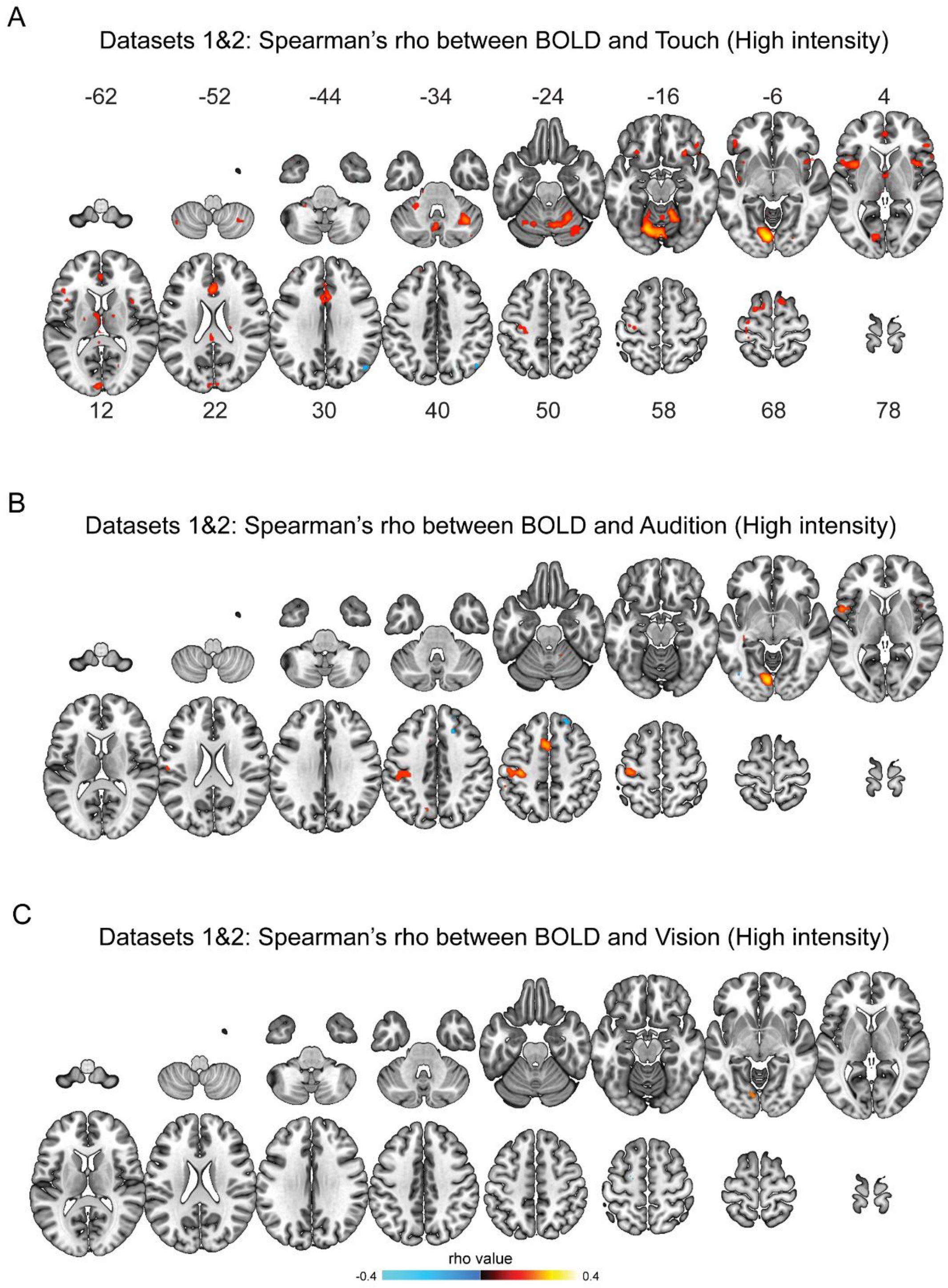


**Figure S5. Robustness of the correlation between BOLD responses and nonpain sensitivity in the high intensity condition in Datasets 1&2.** Nonparametric Spearman correlation between BOLD responses and tactile (A), auditory (B), and visual (C) sensitivity in the high intensity condition in Datasets 1&2. Nonparametric correlation revealed similar correlational patterns as parametric correlation.

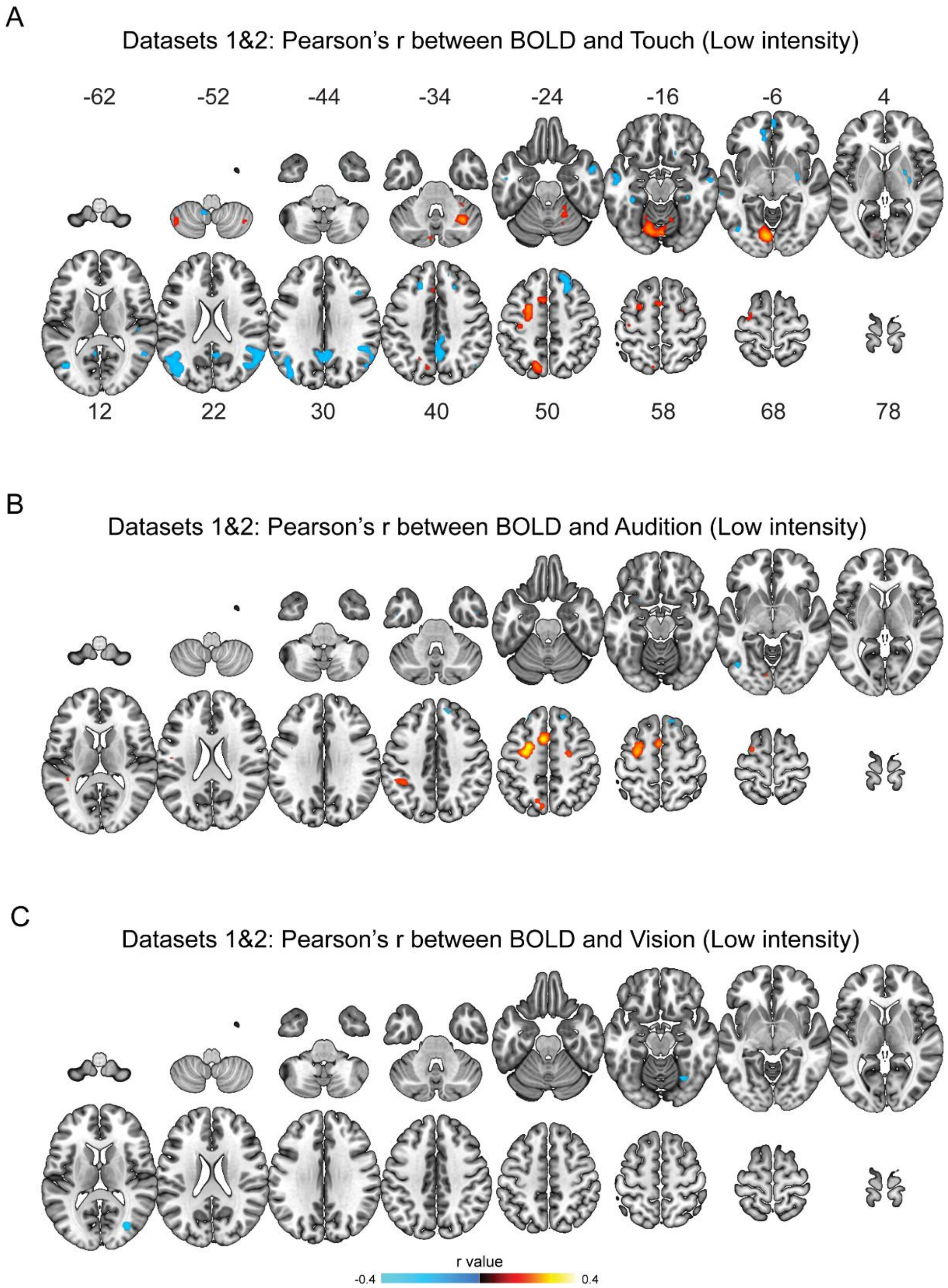


**Figure S6. Correlation between BOLD responses and nonpain sensitivity in the low intensity condition in Datasets 1&2.** Pearson correlation between BOLD responses and tactile (A), auditory (B), and visual (C) sensitivity in the low intensity condition in Datasets 1&2. Correlational patterns in the low intensity condition differed from those in the high intensity condition, suggesting an influence of stimulus intensity.

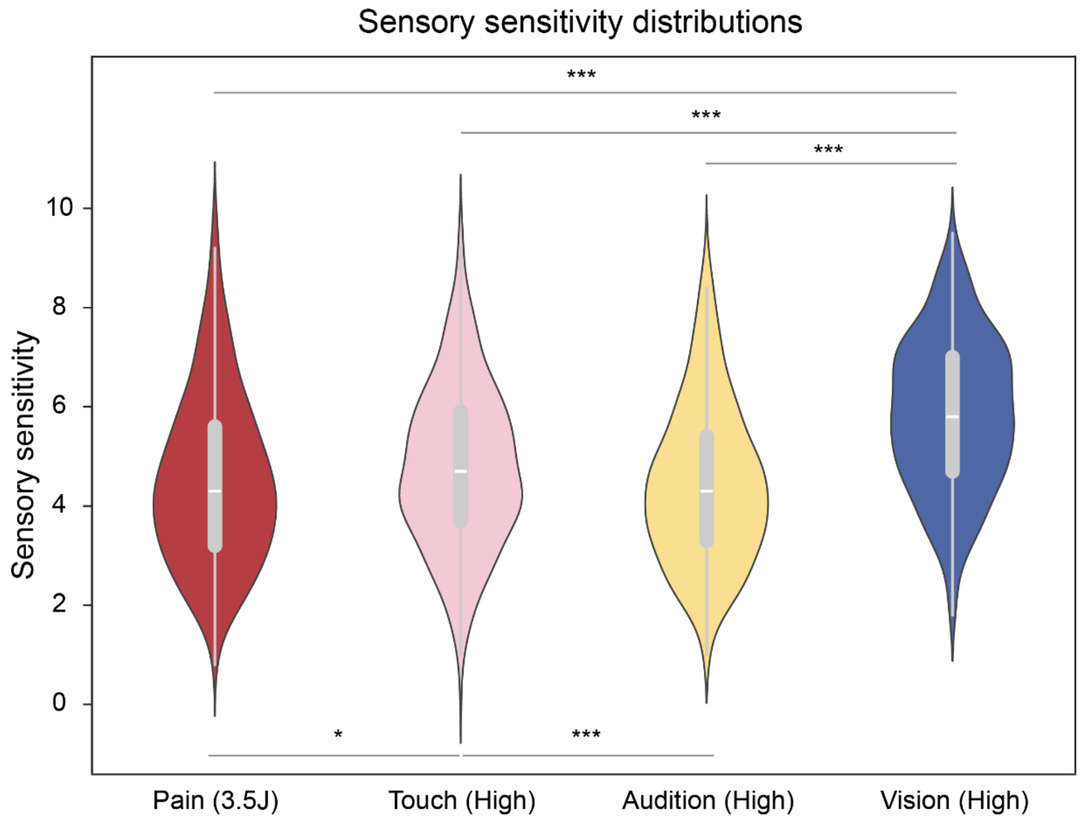


**Figure S7 Distributions of sensory sensitivity in Datasets 1&2.** Sensitivity differed between modalities. Tactile sensitivity was larger than pain and auditory sensitivity, and visual sensitivity was larger than pain, tactile, and auditory sensitivity.

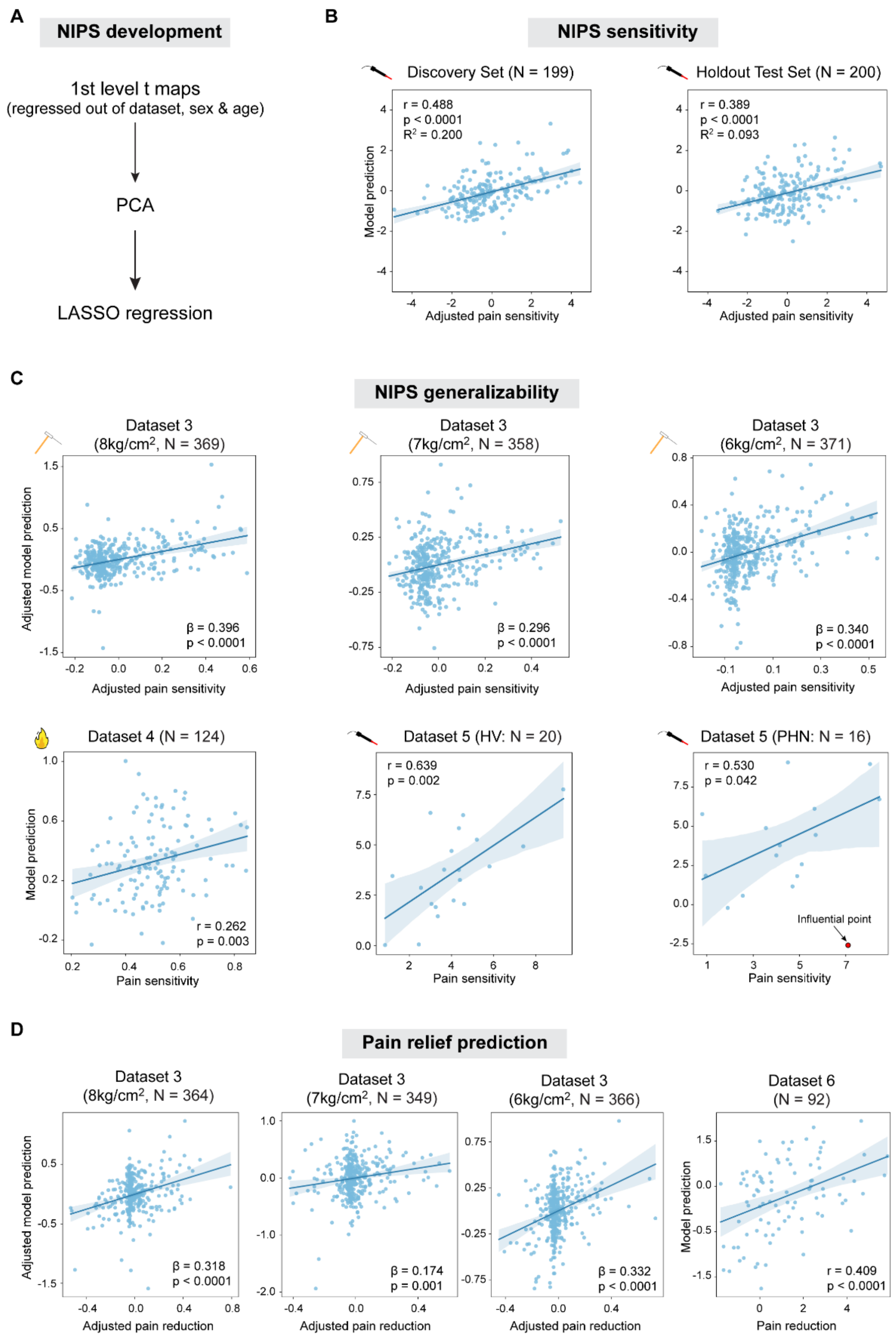


**Figure S8. Performance of multivariate model after regressing out control variables.** The model was built with fMRI responses and pain ratings that were adjusted by regressing out dataset, sex, and age. In panel B (Datasets 1&2), "adjusted pain sensitivity" means that average pain ratings were adjusted for dataset, sex, and age. In panels C&D (Dataset 3), "adjusted" means that fixed effects of sex and age, and random effects of family were adjusted. β in panels C&D is the standardized coefficient in mixed effects models. Note that real and predicted values in Datasets 4~6 were not adjusted since dataset heterogeneity and dependent observations were not present in these datasets. Influential points in Dataset 5 were identified according to Cook's distance > 0.5, and removed to obtain robust results. HV: healthy volunteers; PHN: postherpetic neuralgia patients.

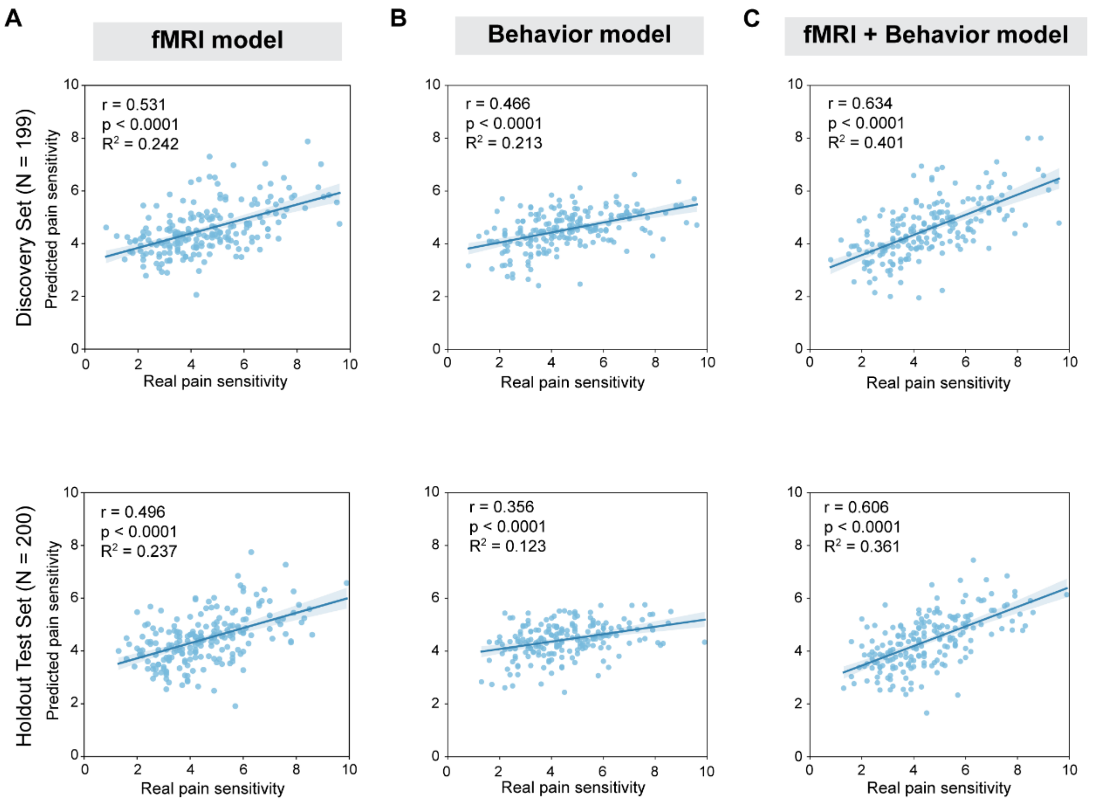


**Figure S9. Performance of fMRI only, behavior only, and fMRI + behavior models.** Compared with fMRI only (A) and behavior only (B) models, a composite model including both fMRI responses and behavioral measures had better performance.

### Supplemental tables

**Table S1. Dataset summary**

| | **Dataset 1** | **Dataset 2** | **Dataset 3** | **Dataset 4** | **Dataset 5 (Patient)** | **Dataset 5 (Healthy)** | **Dataset 6** |
|---|---|---|---|---|---|---|---|
| N | 212[*] | 187[*] | 395 | 124 | 16 | 20 | 92 |
| Sex (M:F) | 76:135 | 103:83 | 164:231 | 63:61 | 5:11 | 8:12 | 42:50 |
| Age (years) | 21.5 ± 4.2 | 21.0 ± 3.3 | 35.4 ± 2.6 | 22.2 ± 2.7 | 65.8 ± 7.0 | 61.6 ± 8.2 | 21.9 ± 3.2 |
| Intervention | / | / | Placebo, control | / | / | / | c-TENS, a-TENS, sham |
| Pain stimulus | Laser heat | Laser heat | Mechanical, contact heat[#] | Contact heat | Laser heat | Laser heat | Contact heat |
| Nonpain stimulus | Electro-tactile, auditory, visual | Electro-tactile, auditory, visual | / | / | / | / | / |
| Intensity level | 2 | 2 | 3 | 6[&] | 1 | 1 | 1[$] |

Note: [*]: One participant did not provide demographic information. [#]: Contact heat pain data were not analyzed. [&]: Data were only available for one intensity level. [$]: Perceived intensity ratings were fixed, not physical intensity.

**Table S2. Functional MRI acquisition parameters**

| Parameters | Datasets 1&2 | Datasets 3&4 | Dataset 5 | Dataset 6 |
|---|---|---|---|---|
| MRI scanner | GE MR 750 | Siemens Prisma | GE MR 750 | Siemens Prisma |
| Magnetic strength | 3T | 3T | 3T | 3T |
| Field of view (mm) | 192 | 220 | 220 | 192 |
| Number of slices | 43 | 56 | 43 | 70 |
| Slice thickness (mm) | 3 | 2.7 | 3 | 4 |
| TR (ms) | 2000 | 460 | 2000 | 2680 |
| TE (ms) | 29 | 27.2 | 30 | 27 |
| Flip angle (deg) | 90 | 44 | 70 | 80 |
| Slice order | Ascending (interleaved) | Ascending (interleaved) | Ascending (interleaved) | Ascending (interleaved) |

**Table S3. Structural MRI acquisition parameters**

| Parameters | Datasets 1&2 | Datasets 3 | Datasets 4 | Dataset 5 | Dataset 6 |
|---|---|---|---|---|---|
| MRI scanner | GE MR 750 | Siemens Prisma | Siemens Prisma | GE MR 750 | Siemens Prisma |
| Magnetic strength | 3T | 3T | 3T | 3T | 3T |
| Field of view (mm) | 256 | 256 | 256 | 256 | 288 |
| Slice thickness (mm) | 1 | 0.8 | 0.7 | 1 | 1 |
| TR (ms) | 6.896 | 2000 | 2400 | 6.9 | 2300 |
| TE (ms) | 2.99 | 2.11 | 2.34 | 2.9 | 2.28 |
| Flip angle (deg) | 8 | 8 | 8 | 8 | 8 |
| Slice order | Ascending (interleaved) | Ascending (interleaved) | Ascending (interleaved) | Ascending (interleaved) | Ascending (interleaved) |

**Table S4. Intensity ratings in different datasets**

| Datasets | Condition | Mean | SD | Max | Min |
|---|---|---|---|---|---|
| Dataset 1 | Pain (3.5J) | 5.294 | 1.637 | 9.900 | 0.800 |
| | Pain (3.0J) | 4.263 | 1.534 | 8.500 | 0.600 |
| | Touch (High) | 4.943 | 1.524 | 9.300 | 1.200 |
| | Touch (Low) | 3.846 | 1.577 | 8.800 | 0.800 |
| | Audition (High) | 4.614 | 1.544 | 9.300 | 1.900 |
| | Audition (Low) | 2.873 | 1.217 | 7.700 | 0.700 |
| | Vision (High) | 6.030 | 1.512 | 9.500 | 2.300 |
| | Vision (Low) | 3.463 | 1.128 | 7.400 | 1.100 |
| Dataset 2 | Pain (4.0J) | 4.547 | 1.480 | 8.800 | 1.200 |
| | Pain (3.5J) | 3.675 | 1.367 | 8.500 | 1.200 |
| | Touch (High) | 4.572 | 1.718 | 9.700 | 0.200 |
| | Touch (Low) | 3.519 | 1.591 | 7.800 | 0.200 |
| | Audition (High) | 4.235 | 1.646 | 8.700 | 1.000 |
| | Audition (Low) | 2.672 | 1.323 | 7.100 | 0.400 |
| | Vision (High) | 5.627 | 1.525 | 9.300 | 1.800 |
| | Vision (Low) | 3.093 | 1.082 | 7.100 | 1.100 |
| Dataset 3 | Control (8kg/cm$^2$) | 0.150 | 0.193 | 0.900 | 0.000 |
| | Control (7kg/cm$^2$) | 0.131 | 0.175 | 0.873 | 0.000 |
| | Control (6kg/cm$^2$) | 0.123 | 0.160 | 0.900 | 0.000 |
| | Placebo (8kg/cm$^2$) | 0.113 | 0.175 | 0.886 | 0.000 |
| | Placebo (7kg/cm$^2$) | 0.092 | 1.148 | 0.816 | 0.000 |
| | Placebo (6kg/cm$^2$) | 0.085 | 0.129 | 0.826 | 0.000 |
| Dataset 4 | Pain (47.5°C) | 0.500 | 0.130 | 0.847 | 0.204 |
| Dataset 5 | Patient (3.5J) | 4.503 | 2.257 | 8.450 | 0.800 |
| | Healthy (3.5J) | 4.070 | 1.927 | 9.300 | 0.850 |
| Dataset 6 | Pre-treatment (Pain) | 6.591 | 1.045 | 9.130 | 4.530 |
| | Post-treatment (Pain) | 5.431 | 1.869 | 9.070 | 1.270 |

### Table S5. Correlation between pain activation and pain ratings in ROIs in Datasets 1&2

| Datasets | ROIs | r | P(raw) | P(FDR) |
|---|---|---|---|---|
| Datasets 1&2 (3.5J) | S1 | 0.248 | <0.001 | <0.001 |
| | S2 | 0.273 | <0.001 | <0.001 |
| | ACC | 0.241 | <0.001 | <0.001 |
| | insula | 0.190 | <0.001 | <0.001 |
| | thalamus | 0.222 | <0.001 | <0.001 |
| Dataset 1 (3.0J) | S1 | 0.194 | 0.005 | 0.005 |
| | S2 | 0.306 | <0.001 | <0.001 |
| | ACC | 0.220 | 0.001 | 0.002 |
| | insula | 0.253 | <0.001 | <0.001 |
| | thalamus | 0.270 | <0.001 | <0.001 |
| Dataset 2 (4.0J) | S1 | 0.211 | 0.004 | 0.009 |
| | S2 | 0.228 | 0.002 | 0.008 |
| | ACC | 0.139 | 0.058 | 0.058 |
| | insula | 0.203 | 0.005 | 0.009 |
| | thalamus | 0.141 | 0.054 | 0.058 |

Note: Cells highlighted in red show significant correlation after FDR correction.

**Table S6. Correlation between pain activation and pain ratings in ROIs in Dataset 3**

| Conditions | ROIs | β | P(raw) | P(FDR) |
|---|---|---|---|---|
| $8kg/cm^2$ | S1 | 0.151 | 0.003 | 0.004 |
| | S2 | 0.149 | 0.004 | 0.004 |
| | ACC | 0.230 | <0.001 | <0.001 |
| | insula | 0.284 | <0.001 | <0.001 |
| | thalamus | 0.149 | 0.003 | 0.004 |
| $7kg/cm^2$ | S1 | 0.080 | 0.121 | 0.121 |
| | S2 | 0.118 | 0.023 | 0.028 |
| | ACC | 0.142 | 0.006 | 0.015 |
| | insula | 0.189 | <0.001 | 0.001 |
| | thalamus | 0.123 | 0.016 | 0.027 |
| $6kg/cm^2$ | S1 | 0.025 | 0.619 | 0.691 |
| | S2 | 0.051 | 0.319 | 0.532 |
| | ACC | 0.080 | 0.117 | 0.293 |
| | insula | 0.104 | 0.039 | 0.197 |
| | thalamus | 0.020 | 0.691 | 0.691 |

Note: β is the standardized coefficient in mixed effects models that take into account observation dependence within families. Cells highlighted in red show significant coefficients after FDR correction.

**Table S7. Post-hoc comparisons between sensory sensitivity in Datasets 1&2**

| Comparisons | | Mean difference | SE | t | df | P(FDR) |
|---|---|---|---|---|---|---|
| Condition1 | Condition2 | | | | | |
| Pain | Touch | -0.234 | 0.091 | -2.573 | 398 | 0.017 |
| Pain | Audition | 0.099 | 0.094 | 1.056 | 398 | 0.291 |
| Pain | Vision | -1.306 | 0.095 | -13.804 | 398 | <0.0001 |
| Touch | Audition | 0.333 | 0.076 | 4.408 | 398 | <0.0001 |
| Touch | Vision | -1.702 | 0.073 | -14.666 | 398 | <0.0001 |
| Audition | Vision | -1.405 | 0.075 | -18.737 | 398 | <0.0001 |
